# A population-scale map of human tandem repeat composition and mutation dynamics from long-read assemblies

**DOI:** 10.1101/2024.08.07.607105

**Authors:** Bida Gu, Dandan Peng, Christy W. LaFlamme, Mark F. Bennett, Melanie Bahlo, Ben Weisburd, Heather Mefford, The Human Genome Structural Variation Consortium, The Human Pangenome Reference Consortium, Mark J.P. Chaisson

## Abstract

Tandem repeats (TRs) are highly mutable genomic elements composed of repeated motifs that drive genetic diversity and contribute to disease. Due to limited resolution at population scale, their mutation dynamics are incompletely understood. Here, we present TRCompDB, a reference database constructed on 832 human genomes that annotates the compositional diversity and population structure for 4.4 million TR loci. Leveraging SNP-based genealogical trees, we estimated locus-specific mutation rates for 4,037,274 TRs, revealing mutation rates ranging from 10^−10^ to 10^−3^ on TRCompDB loci. In addition to TR length and coding potential, mutation rate was influenced by recombination and the wider duplication context. Moreover, a subset of AT-rich pentanucleotide repeats including loci linked to Familial Myoclonic Adult Epilepsy and spinocerebellar ataxia showed extreme mutation rates. Beyond single-motif changes, 29,964 loci exhibited higher-order duplication patterns that increased allelic complexity, among which 3,351 VNTRs showed evolutionary recurrence. These results establish TRCompDB as a resource for studying global patterns of tandem repeat evolution and their contribution to human disease.

## 1 Introduction

Tandem repeats (TRs) are composed of short motif sequences repeated tail-to-head, and comprise 3-4% of the human genome. Loci composed of motifs with fewer than 7 nucleotides are categorized as short tandem repeats (STR) or microsatellites, while those with motifs of ≥7 nucleotides are variable number tandem repeats (VNTRs). While most TR variation is considered benign, advances in algorithmic analysis and long-read sequencing are revealing impacts of TR variation on traits including expression levels, height, and disease risk [1, 2, 3, 4, 5, 6, 7, 8, 9, 10]. TR loci are often studied as a distinct class of DNA because they are subject to multiple mechanisms of mutation including slipped strand mispairing [11], unequal crossover [12, 13], and gene conversion [12, 14], which contribute to a much higher mutation rate and heterogeneity than single-nucleotide substitutions (SNVs). The genetic instability of TR sequences manifests as germline [15] and somatic [16].

Due to associations with molecular and physiological phenotypes, direct impact on disease, and theories that TR sequences serve as rapidly evolving functional elements [17, 18, 19], it is necessary to catalog TR variation and mutation rate in humans. The TR mutation rate has been characterized using direct observations in ex-vivo experiments [20] and as *de novo* variation in pedigrees [15, 21, 22, 23, 24]. These characterized TR variation as a class, for example across all di- and tri-nucleotide repeats. Measuring TR variation across populations enables estimating locus-specific mutation rates given estimates of relatedness between individuals. Population-scale databases of STR variation have been generated for global populations in the Simons Diversity Project [25, 26], the 1000 Genomes Project [27, 28], multiple biobanks [29], and autism [30] using short-read genome sequencing (srGS). Two landmark studies characterized locus-specific TR mutation rates in populations, first on the Y chromosome using directly computed genealogy [31] and subsequently in autosomoal chromosomes using a SNP-based estimation of coalescence [26]. These studies demonstrated that TR mutation rates vary between loci and are up to 10^*−*3^ mutations per generation, roughly five orders of magnitude greater than SNV variation.

Here, we took advantage of three technological advances that enabled more refined estimates of TR mutation rate. First, global reference and pangenome projects construct haplotype-resolved near telomere-to-telomere assemblies of human genomes using long-read genome sequencing (lrGS), where TR variation is sequence-resolved. Second, advances in methods to construct Ancestral Recombination Graphs (ARGs) have improved scalability so that local genealogy may be calculated on hundreds of autosomal chromosomes. Finally, the motif composition of TR sequences may be annotated using our previously developed method, vamos [32], giving global collections of annotations of TR sequences.

The `vamos` method [32] is composed of two modules: one that detects motifs that are repeated at TR loci in a population, and another to annotate TR sequences using these motifs. The efficient motifs are collected from all motifs observed in a population, and filtered to remove motifs with low frequency, balancing similarity to the original motif set and parsimony.

The efforts from pangenome studies have produced long-read sequencing data and high-quality haplotype-resolved assemblies for hundreds of individuals. This provides a valuable resource of sequence-resolved TR sequences (Supplementary Figure S1) [33, 34, 35, 36, 37, 38]. In this study, we applied `vamos` to 416 haplotype-resolved human HiFi assemblies to construct a comprehensive reference of tandem repeat sequence and composition variation, TRCompDB. We also present multiple advances of the `vamos` method and software ecosystem `tryvamos` (Supplementary Notes 1-2) to study tandem repeat variation in large LRS cohorts. Building on prior evidence of strong linkage disequilibrium between STRs and SNPs [39], we estimated used genealogical relationships from SNP data to estimate mutation rates for 4,037,369 autosomal loci (96.8%) in TRCompDB. This provides a global mutational map that will support and inform future research on TRs. Finally, we examined genomic determinants of TR mutation rate and identified candidate TR loci as potential epilepsy-associated targets.

## 2 Results

### 2.1 TRCompDB: an annotation of 4,422,661 TRs among 416 HiFi assemblies

We created a comprehensive catalog of tandem repeats (TRs), TRCompDB, using a *de novo* approach that delineates precise genomic TR boundaries and resolves consistent population motifs using pangenome assemblies. Leveraging 94 haplotype-resolved assemblies from the HPRC Phase 1 dataset and a clustering strategy (Supplementary Note 1), this provides population-level consistency in both repeat boundaries and motif sequences. Using this approach, we also generated motifs for the TRExplorer v1.0 catalog [40] on GRCh38 and further complemented our database, leading to an 3,178,674 increase of loci. The current release of TRCompDB includes 4,422,661 non-centromeric TRs on GRCh38 and 4,155,246 on CHM13, covering 4.2% and 4.0% of the genome, respectively (Figure 1a). The higher number of loci in GRCh38 reflects the imprecise definition of centromeric regions in this reference, which results in the retention of additional centromeric TRs. To provide genomic context on GRCh38, this catalog contains 42,880 TR that overlap with coding sequences as well as 211,019 loci in segmental duplications (SDs) and 1,993,392 TRs in mobile elements (Figure 1b), the latter two categories being historically challenging to profile in previous short-read based databases of TR variation. To assess how well assemblies covered TR loci, we annotated the catalog across all 505 diploid genomes from seven long-read sequencing consortia using `vamos`. Overall, the HPRC Phase 2 assemblies resolved the most TR loci (Figure 1c, Supplementary Figure S2). TR genotyping coverage closely agreed with the overall genome coverage of each assembly (Supplementary Figure S2), confirming robust and consistent representation across datasets. After removing redundant genomes, we included annotations of 416 haplotype-resolved HiFi assemblies in TRCompDB (Figure 1d). Although the catalog encompasses TRs on autosomal and sex chromosomes, we restricted further analysis primarily to autosomal loci due to their distinct evolutionary properties.

**Figure 1:**
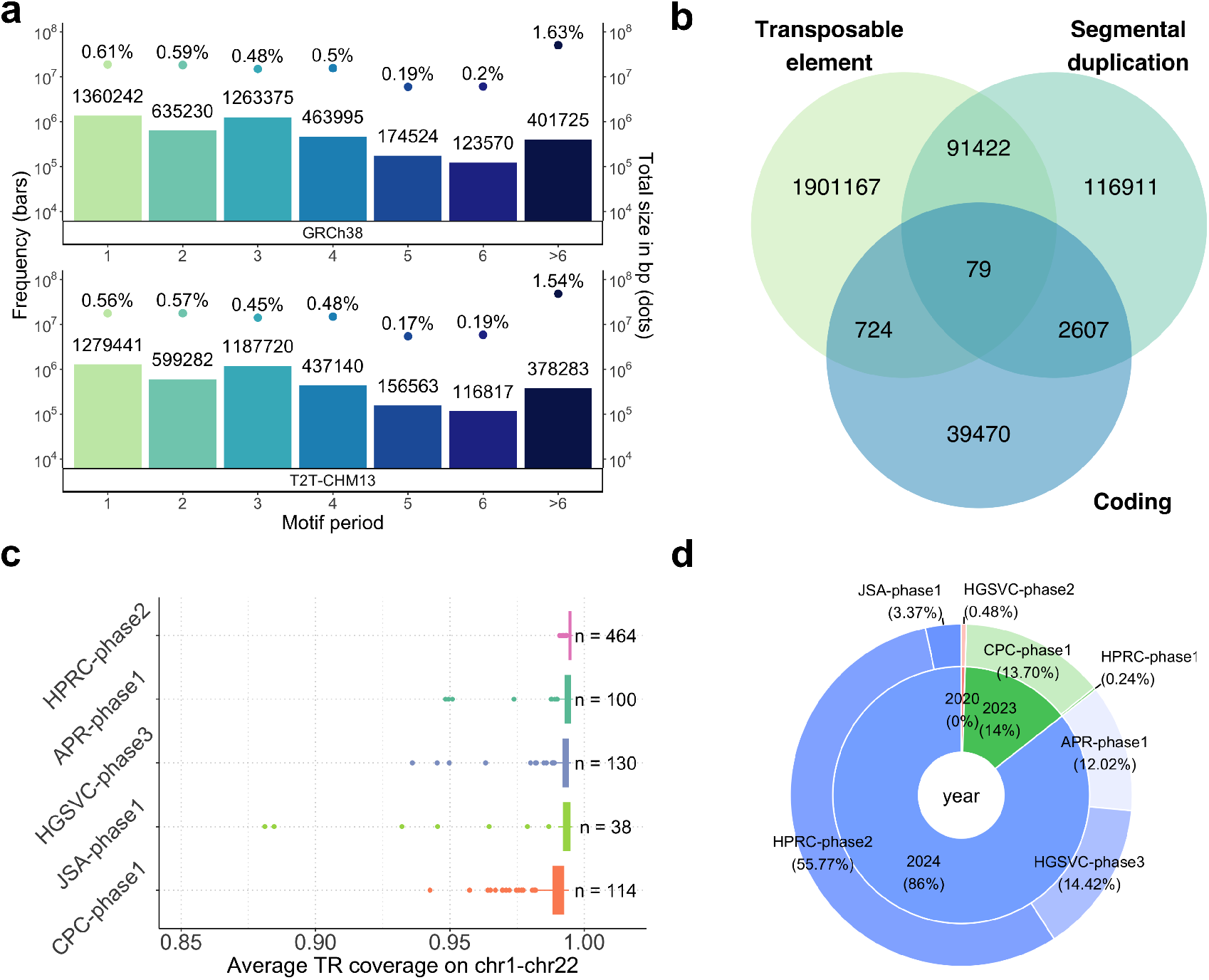
TRCompDB motif catalog and consortium assemblies. **a**, Frequency and size distribution of TRs in the `vamos` databases on GRCh38 and CHM13 reference genomes. bars, TR counts and numbers above; dots, total size and genomic percentages above. **b**, Distribution of TR loci on GRCh38 by overlapping genomic attributes. **c**, TR coverage by `vamos` of genomes in 5 major HiFi sequencing consortia. **d**, Distribution of 416 nonredundant diploid HiFi assemblies in the current TRCompDB.

Our computationally defined TR boundaries contain differences with loci that have canonical disease definitions. For example, the *HTT* STR of `CAG` repeats is biologically recognized to be within `chr4:3074876-3074933`. However, the locus is followed immediately by another STR of `CCG` repeats (Supplementary Figure S3) and are considered a single locus by our pipeline spanning `chr4:3074876-3075052`. To resolve such differences, we manually curated 72 non-centromeric loci that have been reported with biological functions (Supplementary Table 1) [9, 10]. For these loci, including *HTT*, the manually curated boundary and motif definitions are incorporated into the `vamos` annotations and reflect biologically recognized coordinates (Supplementary Figure S4), ensuring that TRCompDB remains consistent with established disease literature.

### 2.2 Population demography of TR variation

Although most TRs show little variability across individuals, TRs are well known to capture ancestry-related signals, and many variable loci have been linked to biological functions. To quantitatively assess TR variability using TRCompDB, we first measured the average number of TR alleles per haplotype-resolved assembly across the four largest consortium datasets on GRCh38, considering both allele length and motif composition (Figure 2a). Consistent with prior expectations, the majority of TR loci were either invariant (60.5% by length, 56.3% by composition) or had an average of < 0.02 alleles per haplotype (37.9% by length, 41.2% by composition) in the population. Despite this overall stability, principal component analysis (PCA) based on either TR allele length or motif composition revealed clear clustering of major population groups (Figure 2b). These clusters closely resemble those observed from SNP data [25], demonstrating that even a subset of variable TRs encodes ancestry information similar to SNP variation.

**Figure 2:**
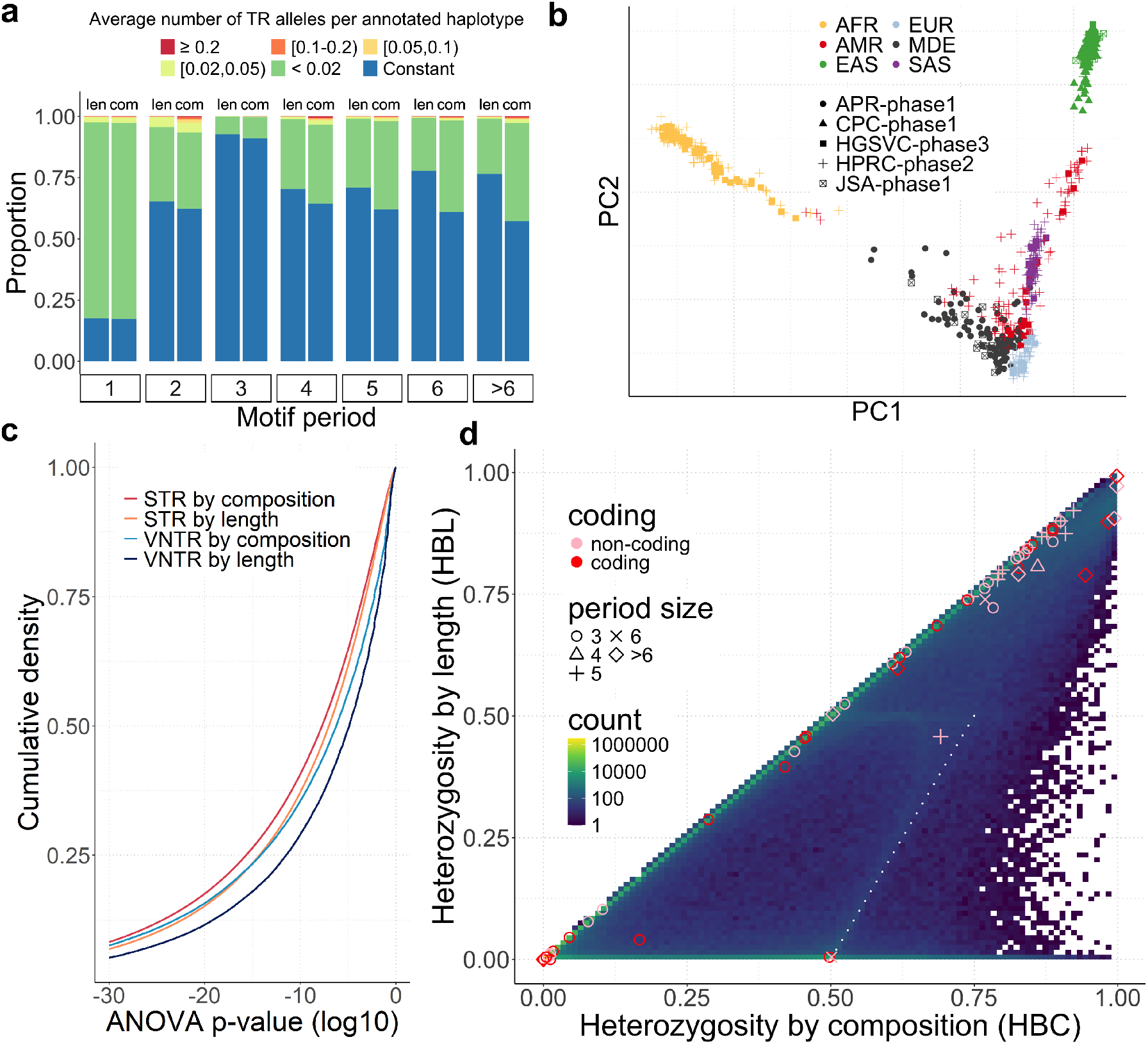
Tandem repeat variation in reference genome cohorts. **a**, TR variation by motif size. For each size group, alleles measured by length are shown on the left, and composition on the right. **b**, Principal component analysis of tandem repeat sequences by population and study. **c**, ANOVA of TR variability within versus between populations, quantifying variation by length and composition differences, shown in order of significance, p-values capped at 10^*−*30^. **d**, Heterozygosity of tandem repeat sequences measured by motif and length variability. Known functional autosomal loci are annotated as affecting coding (N=27, red) or noncoding (N=37, pink) sequences, along with the period size of the disease TR. The while dashed line marks `HBL` = 2`HBC` *−* 1.

We then searched for loci that showed signatures of population differentiation, considering both length and compositional variation. While there is a copy-number (e.g., length) analog to *F*_*ST*_, *V*_*ST*_ [41], this does not extend to multiallelic sequences that vary by composition. To uniformly compare population differentiation by length versus composition, we measured ANOVA according to length variation as well as motif-based edit distance to the major allele for each locus. Considering the 1,136,502 loci that are variable by allele length and have at least 5 annotated genomes in each population, there were 417,681 (36.8%) population informative loci (*p* < 10^−10^). In contrast, motif-based ANOVA revealed an additional 25.8% (525,622) population informative loci (Figure 2c). Moreover, among 780,566 loci that are variable by composition but length, 66,444 additional loci were found to be stratified by population. Population informative TRs were enriched in coding regions when tested by both allele length (*p* = 2.9 ×10^−15^, two-sided, *OR* = 1.4, Fisher’s exact test) and composition (*p* < 2.2 ×10^−16^, two-sided, *OR* = 1.9, Fisher’s exact test). The stronger enrichment effect by the compositional measure further indicates that additional population informativeness can be revealed by compositions. The proportion of population informative STRs is slightly higher than VNTRs (Figure 2c), possibly because STRs are less mutable (Results 2.3) and preserve more population specific alleles. As an example, a coding VNTR sequence *PLIN4* (chr19:4510838-4513560) that is associated with skeletal-muscular disorder [42] was found to have differentiation both by length (*p* = 10^−58^, ANOVA) and motif variability (*p* = 10^−70^, ANOVA; Supplementary Figure S5).

Because allele counts are sensitive to sample size, we next quantified variability at each locus using heterozygosity, defined as 1− homozygosity, where homozygosity is the sum of the squared allele frequencies across all alleles. A completely invariant locus has heterozygosity of 0, whereas highly variable loci exhibit higher values. Loci were characterized by two metrics: heterozygosity by length (HBL) that considers only the length of a locus by motif count, and heterozygosity by composition (HBC) that considers the full motif annotation of a locus. The inclusion of motif composition dramatically increased heterozygosity (Figure 2d). The relative frequency of the second-most frequent allele, akin to a minor allele in biallelic SNP studies, have a much greater frequency than SNP minor alleles, producing the faint enrichment around `HBL` = 0.5. Moreover, additional enrichment along `HBL` = 2`HBC` − 1 (Figure 2d, white dashed line) indicated that incorporating motif composition nearly doubled the apparent heterozygosity at many loci, underscoring the importance of accounting for both length and sequence features in variability analyses. Coding TRs generally showed lower variability than non-coding TRs (6.2-fold change by length, *t* = 186.5, *df* = 45, 848, *p* < 2.2 × 10^−16^, 4.6-fold change by composition, *t* = 162.4, *df* = 44, 644, *p* < 2.2 × 10^−16^, two-sided, t-test), consistent with stronger selective constraint on coding regions [26]. To investigate biological relevance, we highlighted 64 non-centromeric autosomal TRs from the curated list of 72 loci associated with disease or traits. Although most genomic loci exhibited low heterozygosity by composition (< 0.3, 81.7%), the majority of functionally relevant loci were highly variable (≥0.3, 75.0%), and the difference was highly significant (*p* < 2.2 × 10^−16^, two-sided, Fisher’s exact test).

### 2.3 Estimation of locus-specific mutation rates using local ancestry

Given the strong concordance between SNP and TR based ancestry clustering, we next sought to estimate evolutionary mutation rates of TRs by leveraging genealogical information inferred from SNP data and the compositional annotations from `vamos`. We generated an ancestral recombination graph (ARG) using phased SNPs detected from assemblies to estimate local ancestry, and estimated tandem repeat mutation rate using the local tree inferred around each TR locus (Methods 4.1). After applying stringent quality filters for assembly coverage, we computed the mutation rates for 4,037,274 autosomal loci, representing 96.8% of the 4,171,961 loci surveyed. We excluded SNPs inside of TR loci from ARG construction because low-complexity sequences are a source of SNP calling error [43]. The loci excluded from analysis were enriched in subcentromeric regions and chromosomal termini, where genome assemblies are often incomplete.

TRs have been considered to be located in genomic regions with elevated recombination activity [44]. In contrast, we found that the majority of loci in our dataset (3,879,959; 93.0%) are fully contained within a single recombining segment and are thus spanned by a single genealogical tree derived from surrounding SNPs, enabling accurate estimation of evolutionary distances and mutation rates. As shown in Figure 3a, the estimated TR mutation rates span a wide range, from 10^−10^ to 10^−3^ motifs per generation per motif unit. The majority of loci fall within the range of 10^−8^ to 10^−4^, with a peak near 10^−5^. Consistent with prior observations, this confirms that TRs exhibit mutation rates several orders of magnitude higher than the rate of SNVs ([15, 26, 45]).

**Figure 3:**
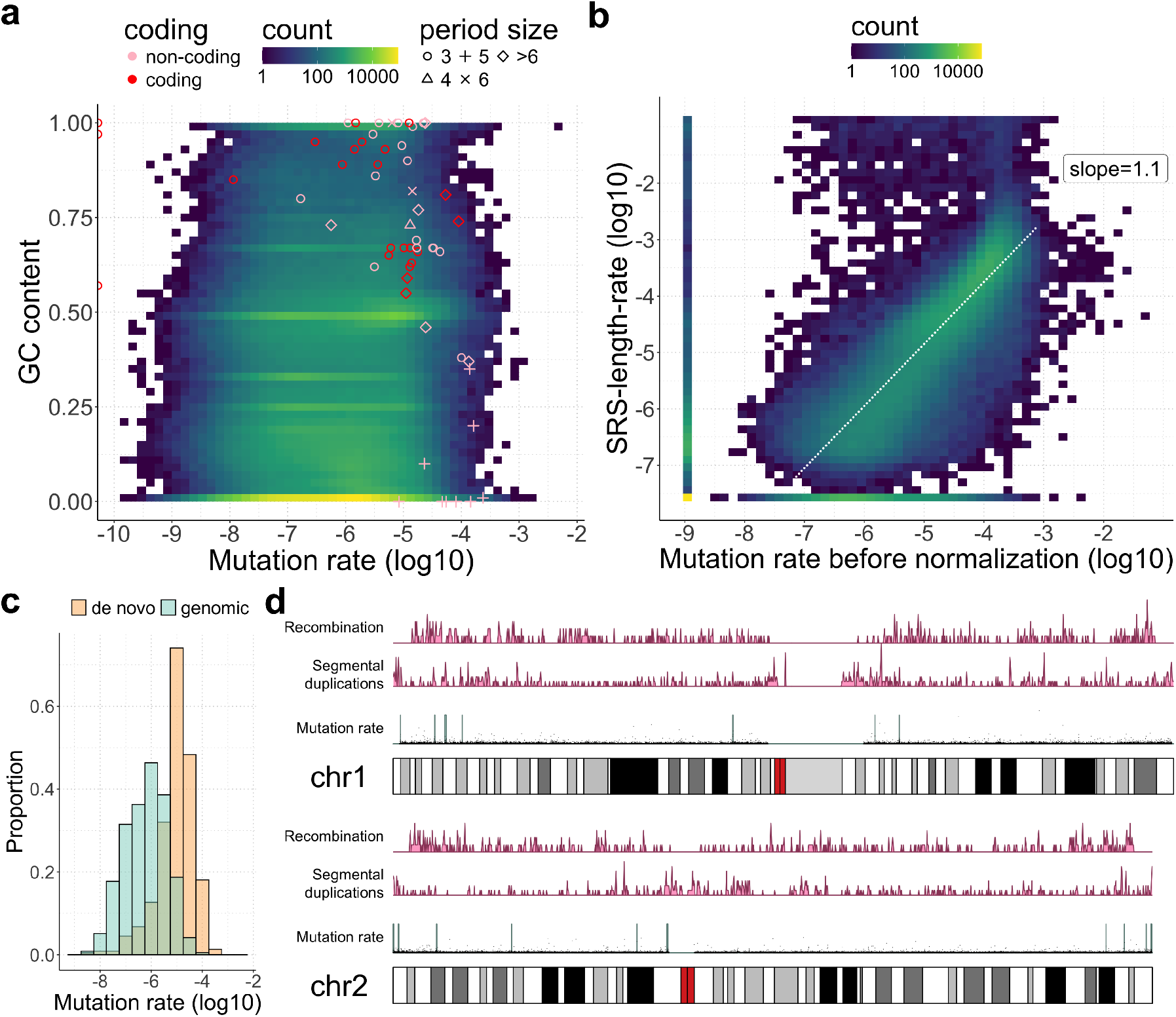
Tandem repeat mutation rate. **a**, Distribution of TR mutation rate, 58 functional TRs are highlighted with corresponding genomic features. The `GC` content reflects overall `GC` composition for all sequences of a TR in the 94 HPRC Phase 1 genomes. **b**, Comparison of TR mutation rate with heterozygosity-based estimates. The mutation rate before normalization by length is compared against previous results. The baseline mutation rate (constant loci) are set at −9 and −7.5 for the two methods. The linear fit line is marked in white. **c**, Mutation rate of *de novo* TRs found in the CEPH1463 study versus genomic TRs. **d**, Segmentation of mutation hotspots on chromosomes 1 and 2. Bottom track, mutation rate (black) and segmented hotspots (green). Middle track, distribution of segmental duplications. Upper track, the distribution of recombination hotspots.

To evaluate the accuracy and robustness of our mutation rate estimates, we compared our results with previously reported STR mutation rates derived solely from allelic length variation (hereafter “SRS-length-rate”) from short-read data[26]. Although both studies employed population genetics–based approaches, several procedural differences exist, including the use of short-read versus long-read sequencing data, distinct TR boundary catalogs, reference SNP mutation rates, sample sizes, and the inclusion of compositional changes in our framework. We lifted the previously reported loci to the GRCh38 reference and restricted analysis to 238,546 loci whose boundaries differed by less than 2 bp between the two catalogs. In addition, we used mutation rates prior to length normalization in our formulation to ensure comparable measurement units. As shown in Figure 3b, despite these methodological differences, the two approaches show high concordance (adjusted *r*^2^ = 0.7, *p* < 2.2 ×10^−16^, linear regression), with the SRS-length-rate estimates roughly 10% higher overall. Nevertheless, three subsets of loci showed systematic discrepancies. First, 11,061 loci (4.6%) were variable in the SRS-length-rate estimates but invariant in ours, likely reflecting the higher accuracy of long-read assembly data compared to short-read sequencing, which tends to overestimate small allelic differences. Second, 23,479 loci (9.8%) were variable in our analysis but at baseline rate in the previous study. Manual inspection revealed that many of these loci are largely invariant across the population but include rare variants in a few genomes. These additional alleles are captured only with our larger dataset (116 more genomes) and the inclusion of compositional changes. Finally, a small subset of 1,324 loci (0.6%) showed markedly inflated SRS-length-rate estimates (> 10^−2.5^) compared to ours, again consistent with genotyping errors in short-read data. Overall, the correlation between the two approaches supports the first orthogonal validation of mutation rate estimates for shared STR loci. Importantly, our long-read assembly–based catalog extends mutation rate estimation to longer and more complex TRs that could not be genotyped using short-read sequencing, including 364,235 VNTRs, of which only 19.5% (71,029 loci) were covered in the previous SRS-length-rate analysis. We also compared our site-specific mutation rates to loci found to have *de novo* mutations in a multi-generation pedigree resolved using lrGS [15]. Of 581 reported loci, 452 were computed with mutation rates in our catalog. The remaining 129 loci were enriched in chromosomal termini that were removed by our coverage quality filter. As expected, the mapped loci shows significantly (*t* = 12.5, *df* = 442.0, *p* < 2.2 ×10^−16^, two-sided, t-test) higher mutation rate than the genomic loci (Figure 3c).

We next explored the relationship between intrinsic TR features and mutation rate. VNTRs are more mutable than STRs (2.9-fold, *t* = 119.9, *df* = 156, 939, *p* < 2.2 × 10^−16^, two-sided, t-test) and loci of low motif purity (< 0.5) are also more mutable (9-fold, *t* = 30.2, *df* = 6, 656.3, *p* < 2.2 × 10^−16^, two-sided, t-test). Here motif purity is defined as 1− impurity, with impurity being the edit distance between a motif to the motif consensus, averaged on all motifs of this locus in the 94 HPRC Phase 1 genomes. There is a strong linear relationship between the log-transformed motif copy number and the log-transformed mutation rate on TRs, for the subset of loci with an average of 5-30 motif copies (adjusted *r*^2^ = 0.2, *p* < 2.2 × 10^−16^, linear regression, Supplementary Figure S6). Regardless of the per-motif normalization, this suggests a power relationship between the number of motif copies and mutation rate for these loci. Nevertheless, we observed statistically significant but weak linear effects on consensus entropy, length, motif purity and locus `GC` content (Table 1), noting that per-motif mutation rate accounts for TR length. This suggests that intrinsic TR features alone cannot fully explain the observed variability in mutation dynamics.

**Table 1:** Association between per-motif TR mutation rate and intrinsic TR features on 1,681,020 variable loci with computed mutation rate. Consensus entropy and length refer to the Shannon entropy and length of the consensus motif sequence. Motif purity is defined as 1*−* impurity, with impurity being the edit distance between a motif to the motif consensus, averaged on all motifs of this locus in the 94 HPRC Phase 1 genomes. Locus `GC` content reflects overall `GC` composition for all sequences of a TR in the 94 HPRC Phase 1 genomes.

| | Adjusted $r^2$ | p-value |
| --- | --- | --- |
| Consensus entropy | 0.02 | $< 2.2 \times 10^{-16}$ |
| Consensus length | 0.03 | $< 2.2 \times 10^{-16}$ |
| Motif purity | 0.01 | $< 2.2 \times 10^{-16}$ |
| Locus GC content | 0.01 | $< 2.2 \times 10^{-16}$ |

### 2.4 TR Mutational hotspots exhibit strong association with local genomic context

To further explore genomic factors that influence TR mutation rates, we analyzed the distribution of highly mutable TRs across the genome. We first assessed whether or not there were “hotspots” of mutation where there was an elevated TR mutation rate relative to the rest of the genome (Methods 4.2). After excluding regions with low SNP density or consistently short SNP tree branch lengths that may be potential artifacts of ARG construction, we identified 95 hotspot regions of elevated mutation rate (Methods 4.2; Figure 3d; Supplementary Figure S7). The hotspots are unlikely to arise at random (*p* < 0.001, permutation test; Methods 4.2), suggesting that local genomic context plays a critical role in shaping TR mutability. We next tested whether these hotspots were enriched for known genomic features. Recombination hotspots [46] were significantly associated with increased TR mutation rates (*p* = 2 ×10^−8^, two-sided, Fisher’s exact test). While this is consistent with the role of recombination as a mutational driver of TR variation, only 0.19% of variance is explained by proximity (within 10kb) to recombination hotspots, suggesting a minor effect. Mutational hotspots were also significantly enriched for non-coding regions (*p* < 2.2 ×10^−16^, two-sided, Fisher’s exact test), consistent with the expectation that reduced selective constraint in these regions allows TR variation to accumulate more readily. Most strikingly, we observed a strong enrichment of TR mutation hotspots within segmental duplications (SDs) (*p* < 2.2 × 10^−16^, two-sided, Fisher’s exact test) where TRs overlapping these regions exhibited, on average, a 2.1-fold increase in mutation rate compared to TRs outside SDs.

The elevated mutation rate in SDs may be caused by misaligned paralogs in complex or structurally variable regions. To test this, we annotated all SD sequences as quiescent, high identity (>99%), or structurally variable/invariable, and further quantified hotspot burden within each category. The identity of duplications were determined as the maximum overall reference annotations of a locus using SEDEF [47]. The quiescent SDs are ancient (low-identity), not mutating sequences that are easily mapped to orthologous GRCh38 loci, while high identity or structurally variable regions may cause misaligned segments during whole-genome alignment [48]. To quantify structural variation, reference SD along with 50kb flanking sequences were extracted and aligned to each assembly to determine the relative length of the orthologous sequence in the assembly. The SD loci with length difference greater than 10% were annotated as structurally variable. An enrichment of hotspot loci was observed in all classes of SDs, although with a greater effect in high identity and structurally variable SDs (Table 2). Moreover, both the low identity and structurally invariant SDs exhibit an increased mutation rate (2.0-fold and 1.9-fold) compared to TRs outside SDs, sufficient to explain that the overall elevation of TR mutation rate by SDs is not a result of misalignment.

**Table 2:** Distribution of TRs in mutation hotspots by classes of segmental duplications.

| Segmental duplication class | hotspot | normal | total |
| --- | --- | --- | --- |
| High identity | 611 (7.4%) | 7,650 (92.6%) | 8,261 |
| Low identity | 2,174 (4.5%) | 45,852 (95.5%) | 48,026 |
| High structural variation | 592 (14.7%) | 3,442 (85.3%) | 4,034 |
| Low structural variation | 2,193 (4.2%) | 50,060 (95.8%) | 52,253 |
| Not in segmental duplications | 4,022 (0.2%) | 1,620,711 (99.8%) | 1,624,733 |

### 2.5 Highly mutable AT-rich pentanucleotide repeats may provide potential target for epilepsy associated tandem repeats

With the constructed genomic TR mutational catalog in the healthy TRCompDB population, we next examined the mutational profile of the 72 manually curated functional TRs, and discovered highly mutable `AT`-rich pentanucleotide STRs as candidate epilepsy associated tandem repeats. Of the 64 non-centromeric autosomal functional TRs, mutation rates were successfully computed for 58 loci (Figure 3a). Overall, the functional loci exhibit medium (10^−6^) to high (10^−4^) mutation rates, suggesting that TRs involved in biological function are less likely to be evolutionarily stable. As expected, non-coding TRs show significantly higher average mutation rates (4.3 × 10^−5^) compared to coding TRs (1.2 ×10^−5^; *t* = 2.9, *df* = 44.2, *p* = 0.006, two-sided, t-test). A small subset of extremely `AT`-rich pentanucleotide repeats shows high mutation rates (> 10^−5^), and are associated with Familial Adult Myoclonus Epilepsy (FAME) and Spinocerebellar Ataxia (SCA) (Supplementary Table 2). Manual inspection revealed that these repeats are intronic and located near Alu elements, which may contribute to their elevated mutation rates. Applying a filter of mutation rate > 10^−5^, `GC` content < 0.05, and repeat length between 40 and 120 bases, we identified 343 extra `AT`-rich pentanucleotide repeats that can be transcribed (Supplementary Table 3). Since most known disease-associated loci of this class also show a high length standard deviation in the population, we further refined the list to 27 loci (Supplementary Table 4), most of which are in close proximity to Alu elements. Outlier test by motif copy number revealed that 26 loci had outlier expansions that were 10 times of the inter-quartile range (IQR) above the 3rd quartile, consistent with the reported disease-associated `AT`-rich pentanucleotide loci (Supplementary Figure S8). In particular, one of the loci (chr1:97316514-97316562; Supplementary Figure S9) was found to be within the DPYD gene from the Genes4epilepsy list [49], potentially highlighting an unstable locus. While none of these loci were found to be expanded among 170 probands from a srGS study on pediatric epilepsy, nor among 15 srGS samples for individuals with FAME or later-onset progressive myoclonic epilepsy [50], their high similarity to known disease-associated loci and elevated mutation rate highlights them as promising novel disease loci.

### 2.6 Higher-order repeat structure of euchromatic TRs

We frequently observed higher-order of repetitive patterns within TRs, which involves the gain (or symmetrically the loss) of multiple motifs between phylogenetically adjacent individuals (Figure 4). Such events may occur in coding TRs and result in substantial allelic differences, as shown by the MUC1 coding VNTR (Supplementary Figure S10). We reasoned that these mutations, referred to as block mutations, arose from unequal crossover events during recombination, or by other mutational mechanisms than polymerase slippage that is predicted to make only small changes to motif copy number [8]. To systematically classify such events, we developed an approach to classify differences between sequences as having arisen through a large “block” duplication encompassing multiple motifs, versus slippage events (Methods 4.3). Across 2,221,932 autosomal TRs with motif sizes greater than two nucleotides and annotated in at least 20 genomes, 29,964 loci (1.3%) were classified as having block mutations (block-positive), indicating that these events constitute a notable component of TR mutational diversity.

**Figure 4:**
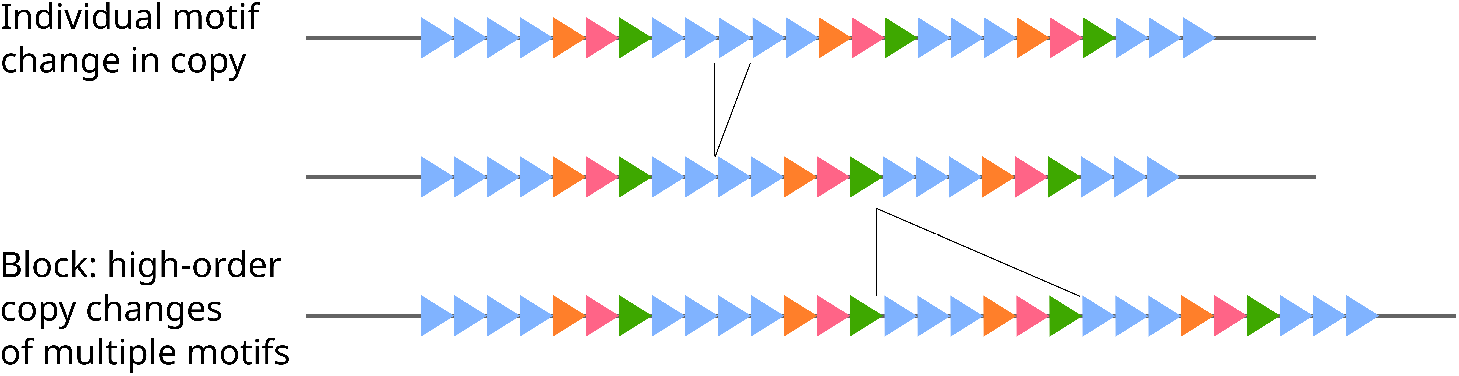
Block mutations. Three TR alleles are shown with motifs represented by triangles. Mutation of a single blue motif is observed between the top and middle allele. Multiple motifs are duplicated from the middle to the bottom allele. This results in higher-order of repeats (of block of motifs) in a TR sequence and substantial allelic differences.

We next examined genomic correlates of these block-level events. In non-telomeric regions, loci without block mutations were on average 1.14 times farther from recombination hotspots than block-positive loci (*t* = 13.2, *df* = 26, 056, *p* < 2.2 × 10^−16^, two-sided, t-test), indicating an association between recombination and block-level mutations. Moreover, block-positive loci exhibited a 5.1-fold increase in the average number of alleles defined by length, highlighting the substantial enrichment of allelic diversity driven by block mutations. Among 4,653 block-positive VNTRs fully spanned by SNP genealogical trees, 3,129 loci (67.2%) contained more than one block pattern cluster. The heterozygosity by length increased with the number of distinct block pattern clusters (adjusted *r*^2^ = 0.1, *p* < 2.2 × 10^−16^, linear regression), indicating that loci harboring multiple block configurations exhibit more complex allelic architectures. Finally, to quantify the recurrence of block mutations, which is not captured by overall mutation rates, we leveraged SNP-based genealogical trees to reconstruct the evolutionary histories of TR alleles for these 4,653 VNTRs using maximum parsimony reconstruction (Methods 4.4), and counted recurrence according to block changes between ancestral sequences. Overall 3,351 (72.2%) loci showed recurrent mutations. As expected, the number of recurrent block mutations increased with the mutation rate (adjusted *r*^2^ = 0.2, *p* < 2.2 × 10^−16^, linear regression). However, no significant relationship was observed between recurrent block mutation frequency and distance to recombination hotspots. Together, these findings demonstrate that block-level mutational processes are both common and they play a major role in expanding TR allelic diversity in human populations.

## 3 Discussion

In this study, we presented a reference catalog for TR variation of over 4 million sites from 832 assembled haplotypes. The use of high-quality pangenome assemblies yields additional insight compared to previous short-read based studies by resolving sequence composition, variation in challenging to map segmental duplications, and regional phasing that enables ancestral reconstruction. For example, including motif diversity gives an average of a 2.4-fold increase of heterozygosity compared to length-based estimates.

The resulting inferred mutation rate serves as orthogonal validation of previous estimates based on short read data [26], largely adding complex loci that cannot be profiled using short-read sequencing. The ARG-based approach to estimate ancestry allows for estimating ancestral states and counting mutation recurrence. Future refinements of TR mutation rate can incorporate a unified model of slippage, block duplications, and recurrence, all leveraging genealogy. While an increase in TR variation is frequently observed in subtelomeric regions of the genome [33], the *de novo* assemblies used to annotate TR variation are often unresolved in these regions. In total, our analysis excludes 25Mb of subtelomeric, non-acrocentric DNA. This category of rapidly evolving sequence, along with more complex structures such as rDNA and centromeric loci will require more sophisticated approaches for genealogy reconstruction and comparison of complex repeat loci, possibly leveraging pangenome references in order to account for alignment challenges in these regions.

The ARG-derived mutation rates may be affected by inaccurate graph estimation at recombination hotspots, or when the mutation distance between individuals is low. Tandem repeats have long been considered as drivers for recombination [12], including at specific minisatellites [44]. However, earlier studies on additional markers have indicated no significant enrichment of mutational burden at sites of recombination [51, 52]. Here, leveraging our genome-wide database of TR mutation, we were able to quantify a statistically significant, but minor effect on the mutation rate of TRs through recombination. Furthermore, most TR loci (93%) are spanned by trees, indicating they are rarely associated with recurrent recombination, and the remaining 3.2% unresolved loci are primarily due to incomplete assemblies, SNP density, and filtering on tree branch lengths. Finally, the effect of short branch-lengths from infrequent SNP variation is limited to 0.6% of analyzed loci where the average branch length was under 10,000 (lowest 0.7%), where there is a limited change in the range of mutation rate (Supplementary Figure S11).

A striking feature of TR variability is its heterogeneity, even within the same repeat motif. When considering all loci with the same STR motif, the majority of STR classes show an over 10-fold difference for the inter-quartile range of mutation rate (Supplementary Figure S12). The most variable STR motif, `AT`, had a 40-fold difference in span within the interquantile range of mutation rates, distributed genome-wide (Supplementary Figure S13). The mutation rates in TRcompDB are normalized per motif, effectively removing the largest known predictor of TR variability, length. Comparing nearly equivalent length loci can emphasize diversity of mutation rates. For example, among `AT` STRs, when limiting to those from reference loci 50-60 bases, the range of mutation rates spans five orders of magnitude. Because the global features analyzed in this study only account for a limited amount of variance of mutation rate, additional studies examining the propensity for a locus to become mutable are warranted.

The TRcompDB serves as a resource for disease studies by annotating potentially genetically unstable loci. The majority of known disease loci have a high mutation rate, and can operate through a mechanism of lifelong mutation until reaching a point of pathogenicity [16], potentially late enough in life to not affect reproductive fitness and reflect constraint. As an example, pentanucleotide repeat expansions associated with FAME may occur somatically [53]. A recent study by the *All of Us* program presented evidence for two novel pathogenic expansion candidates of `CAG` repeat [54], TRcompDB contains 27 `AT`-rich loci found to have similar sequence characteristics and mutation rates as known FAME risk alleles. The presence of outlier repeat expansions at these loci, their overlap with known epilepsy-associated gene, and possible pleiotropy, highlight them as potential novel disease-associated candidates. More widely, the size of the pangenome allowed the discovery of 19,757 loci with rare and extreme expansions (>10 ×IQR above the 3rd quartile, minimum repeat length standard deviation of five), reflecting genetically unstable loci whose functional impact warrants further investigation.

Our study has several limitations. First, TR genotyping was performed using the efficient motif sets defined by `vamos`, which may underestimate variability and mutation rates due to the exclusion of rare motifs. This tradeoff substantially reduces sequencing noise, and we chose a mild motif replacement threshold (*q* = 0.1) to balance sensitivity and computational feasibility. Second, our mutation rate estimates are based on population genetic inference using SNP-derived local ancestry, which provides an indirect measure of TR mutation dynamics. While pedigree-based approaches can yield more direct estimates, such data remain limited and costly to obtain. The CEPH1463 family study [15] reported an average of 65.3 *de novo* TRs per sample and estimated a genome-wide TR mutation rate of 4.74 × 10^−6^. Our results revealed 98,863 loci whose mutation rates are above 10^−5^, which would require at least 1,514 family-based samples to be directly observed. Therefore, our population-scale framework offers a valuable and comprehensive view of the genomic landscape of TR mutability. Finally, our catalog construction pipeline relies on TRF and RepeatMasker annotations, which include only repeats exceeding minimum alignment thresholds. This limits the detection of shorter TRs, resulting in missing loci in our catalog. Nonetheless, we supplemented these loci by incorporating the TRExplorer v1.0 catalog and computing corresponding motifs within our infrastructure.

Moreover, because our pipeline leverages a pan-genome framework, short loci missed by this threshold are likely to exhibit limited population variability.

In summary, our work provides the most extensive genome-wide analysis to date of human TR variability and mutation dynamics. By integrating high-resolution motif-level annotations, population-scale data, and SNP-based genealogical reconstruction, we reveal how local genomic context and higher-order mutational processes shape TR diversity. These findings not only advance our understanding of TR evolution but also offer a valuable resource for investigating the roles of tandem repeats in human disease and genome function.

## 4 Methods

### 4.1 Calculation of TR mutation rate

TRs were annotated by vamos (v-2.2.0) [32] in the `–contig` mode under the default settings. Genomic SNPs were genotyped by Dipcall (v-0.3) [55] under the default settings, followed by construction of ARGs using Relate (v-1.2.1) [56] with the command `[–mode All -m 1.25e-8 -N 30000 –seed 1]`. To ensure high accuracy, SNPs inside centromere, TRs, or not covered in any genomes were filtered. Constant SNPs across all genomes are not informative and thus also filtered. To cope with samples with large uncovered regions, we partitioned the genome into 2Mb windows and for each window removed samples missing more than 100kb. An ARG was then inferred for every genomic window from the specific set of well-covered samples, resolving the window into genealogical trees, each spanning a recombining segment as inferred from the SNP data. Given the abnormally reduced branch length on trees spanning large genomic regions or with low SNP density, we further removed trees larger than 10kb or having fewer than 1 SNP for every 500bp on average. For a given TR locus, the mutation rate by any pair of haplotypes was calculated as the ratio between the pairwise edit distance of the motif annotation strings to the pairwise branch length (in generation unit) inferred from the tree fully spanning this locus. The overall mutation rate of a locus was obtained by averaging over all pair of haplotypes. For loci not spanned by a SNP tree, if the closest nearby tree is within 10kb, a rescued mutation rate was estimated using this tree. Finally, to account for elevated mutation rate at longer loci, locus mutation rate was normalized by the average motif copy number. The resulted normalized mutation rate has a unit of number of motif changes per generation per motif. Analysis was based on the per-motif mutation rate unless otherwise specified.

### 4.2 Segmentation and permutation test of TR mutation hotspots

Variable TRs with estimated mutation rate were first grouped into bins of 20 loci. Hotspot tags were assigned if the average TR mutation rate of the bin is 1 standard deviation above the chromosomal mean mutation rate. If two hotspot bins were separated by no more than 4 bins, the entire region was merged into a single hotspot. Hotspot bins that did not undergo any merging were subsequently treated as noise calls, whose tags were flipped back to normal. Finally, hotspot regions whose average tree branch lengths are 1 standard deviation below the chromosomal average were filtered. Permutation tests were performed for 1,000 runs, to shuffle the hotspot tags of individual bins before subsequent merging and filtering steps.

### 4.3 Identification of block duplications

For one motif annotation string, all *k*-mers of size 3 or greater were first collected. Since adjacent duplication blocks have matching *k*-mers in their corresponding positions, adjacent matching *k*-mers may signal potential duplication blocks. Therefore, distance between the two matching *k*-mers gives size of the potential duplication block and extending each kmer accordingly by the pattern size gives the potential duplication block sequence. To avoid single duplication event, *k*-mers occurring less than 3 times in the annotation string were not considered. In addition, to avoid homopolymer and nested duplications, cases with potential pattern sizes smaller than 3 were also filtered. Next, if the edit distance between the two candidate patterns is smaller than 20% of the duplication size, both patterns were recorded as candidate duplication pattern. Given a TR locus with at least 20 annotated genomes, if 10 or more genomes were observed with candidate duplication pattern, the locus was tagged as block duplication positive. Finally, we require that candidate patterns be grouped by similarity to give the final list of independent block duplication patterns. This was achieved by checking the cyclic rotations of every candidate pattern against other candidate patterns. Patterns were grouped into the same similarity cluster when the edit distance of any cyclic rotations between them is smaller than 40% of the pattern size. However, such check over all cyclic rotations is computationally expensive and block duplication of STR motifs may simply be interpreted as VNTRs. To reduce the overall computational burden, we only applied the grouping step to VNTRs that were fully spanned by SNP trees.

### 4.4 Inference of TR mutational history to reveal recurrent mutational events

While mutation rates capture the overall mutability of a TR locus over time, they do not fully reveal the underlying mutational dynamics. Moreover, hierarchical clustering based on TR allele distances tends to group alleles by sequence similarity, which can obscure recurrent mutational events. To overcome these limitations, we leveraged SNP-based genealogical trees to reconstruct the evolutionary history of TR alleles. Unlike TR-based clustering, SNP-inferred phylogenies provide an independent framework that is not biased by TR sequence similarity, making them well suited for identifying recurrent TR mutations. In this framework, genomes are represented as leaf nodes. For a specific pattern cluster of a locus, given the observed number of block mutations of each genome, inferring ancestral number of block mutations reduces to estimating the states of internal tree nodes under an optimality criterion. This problem is formally defined as the Small Parsimony Problem. Using the Sankoff algorithm [57], we reconstructed the historical allelic states for block duplication counts across block pattern clusters in all block-positive VNTRs that were fully spanned by SNP genealogies. The total number of recurrent block mutations of a locus was calculated as the sum of recurrent block mutations of all pattern clusters of the locus. To make counts comparable across the genome, we further normalized the count by the average number of motifs in each allele.

### 4.5 Statistical tests

Population-level ANOVA were performed independently at each TR locus using one-way ANOVA in `Python 3.10.14` with the `f_oneway` function from `SciPy 1.14.1` [58]. The corresponding F statistics and degrees of freedom for all locus-specific tests were uploaded to the Zenodo repository (see Data Availability). All two-sample t tests were conducted in `R 4.4.2` using the `t.test` function. Linear regression analyses were performed in `R 4.4.2` using the `lm` function. Fisher’s exact tests were also carried out in `R 4.4.2` using the `fisher.test` function.

## 5 Competing interests

No competing interest is declared.

## 6 Author contributions

B.G. wrote the manuscript and conducted the analysis. M.J.P.C. wrote the manuscript. D.P. helped on running the ARG construction tools. C.W.L, Ma.B., Me.B., H.M. conducted FAME repeat analysis. B.W. contributed TRExplorer loci.

## 7 Funding

B.G. and M.J.P.C were funded by R01HG011649 and U24 HG007497. D.P. was funded by R35GM137758.

C.W.L. has been funded through the St. Jude Graduate School of Biomedical Sciences and the National Institutes of Health (NIH) National Human Genome Research Institute (NHGRI) F99/K00 Fellowship (1F99HG014072). This work was possible by Victorian State Government Operational Infrastructure Support and Australian Government NHMRC IRIISS. Me. B was funded by an Australian National Health and Medical Research Council Investigator Grant (APP1195236).

## 8 Data availability

- All motif catalogs, vcf of TR annotations (TRCompDB), locus-specific ANOVA results, and mutation rate results are available through Zenodo at https://zenodo.org/records/13263614.
- The Human Genome Structural Variation Consortium (HGSVC) Phase 2 assemblies are available through http://ftp.1000genomes.ebi.ac.uk/vol1/ftp/data_collections/HGSVC2/release/v1.0/assemblies/.
- The Human Genome Structural Variation Consortium (HGSVC) Phase 3 assemblies are available through https://ftp.1000genomes.ebi.ac.uk/vol1/ftp/data_collections/HGSVC3/working/.
- The Human Pangenome Reference Consortium (HPRC) Phase 1 assemblies are available through https://github.com/human-pangenomics/HPP_Year1_Assemblies/blob/main/assembly_index/Year1_assemblies_v2_genbank.index.
- The Human Pangenome Reference Consortium (HPRC) Phase 2 assemblies are available through https://humanpangenome.org/hprc-data-release-2/.
- The Chinese Pangenome Consortium (CPC) Phase 1 assemblies are available upon request through https://ngdc.cncb.ac.cn/bioproject/browse/PRJCA011422.
- The Arab Pangenome Reference (APR) project Phase 1 assemblies are available through https://www.mbru.ac.ae/the-arab-pangenome-reference/.
- The Japan and Saudi Arabia (JSA) Pangenome Graph Phase 1 assemblies are available through https://www.ncbi.nlm.nih.gov/bioproject/PRJDB19680/.

## 9 Code availability

- The catalog construction pipeline is available at https://github.com/ChaissonLab/vamos/tree/master/buildTRs.
- `tryvamos` is available at https://github.com/ChaissonLab/vamos/tree/master/tryvamos.
- All code for analysis is available at https://github.com/ChaissonLab/vamos/tree/master/analysis.

## Notes

### Competing Interest Statement

The authors have declared no competing interest.

### Summary of Updates

The manuscript was updated to include new analysis on the tandem repeat mutation rate; More long-read assemblies and tandem repeat loci were incorporated into the database; added new contributing authors.

