## Supplementary Information for "A population-scale map of human tandem repeat composition and mutation dynamics from long-read assemblies"

### Supplementary Notes

#### 1. Construction of TR boundaries and motifs from HPRC Phase 1 genomes

We created our TR locus and motif database using 94 HPRC Phase 1 haplotypes [[1]](https://paperpile.com/c/CapcoG/VUc5). Tandem repeats were first annotated using Tandem Repeat Finder v4.09.1 [[2]](https://paperpile.com/c/CapcoG/MaFr) and RepeatMasker 4.1.2-p1 [[3]](https://paperpile.com/c/CapcoG/73Ym) with default parameters. Each TR call was characterized by an interval in the genome and a consensus motif that was repeated, potentially with variation, within this interval. The TR calls sharing overlaps on the same assembly were merged. In the majority of instances the shortest consensus motif among all merged calls was selected as the new consensus motif. In relatively few instances (< 5%), short TRs were nested in higher-order TRs, and the annotation of the larger instance more accurately reflected the tandem repeat structure of the locus. To address this, we defined nested repeats as regions where the TR call by the shortest consensus motif spanned less than half of the entire region of all overlapping calls. In these instances the motif for the longest overlapping annotation was selected if it resulted in more consistent motif decompositions. Subsequently, TRs with merged boundaries in each genome were mapped to the GRCh38 and CHM13 reference genomes, ready for subsequent processing across genomes.

Constructing a TR database from multiple genomes is challenging due to inconsistencies in TR boundaries and motif decomposition. A TR with a 3-base consensus in one genome might be misinterpreted as a 6-base consensus in another, leading to discrepancies and complicating allele comparisons. This is not explicitly solved by vamos efficient motif selection. To address this, we used k-means clustering to identify inconsistent TR boundaries and motifs before merging TR boundaries across genomes. The clustering groups overlapping TR entries based on genomic proximity and consensus similarity, using boundary coordinates and consensus motif lengths as input features (Supplementary Figure S14). The optimal number of clusters was determined by the highest Silhouette score. Clusters supported by fewer than three genomes were excluded. Each remaining cluster was refined by averaging boundary coordinates and deriving a new motif consensus sequence from the multiple sequence alignment of all motif consensus sequences within the cluster. Overlapping clusters were merged into a single TR locus, with the motif consensus from the most-supported cluster used to re-decompose each genomic TR sequence into motifs via the StringDecomposer algorithm [[4]](https://paperpile.com/c/CapcoG/LpaG). To ensure proper decomposition, boundary motifs differing by more than 20% in length from the consensus, as well as single-base motifs for non-homopolymers, were excluded. The remaining motifs formed the complete motif set for each locus. For computational efficiency, TR loci longer than 10,000 bases or with more than 500 unique motifs were filtered out. Finally, efficient motifs were obtained with a compression level of q=0.1 as described previously [[5]](https://paperpile.com/c/CapcoG/ELYS).

To incorporate extra loci from the TRExplorer v1.0 GRCh38 catalog [[6]](https://paperpile.com/c/CapcoG/OTvl), we first merged overlapping TRExplorer v1.0 loci into clusters and selected the consensus motif of the longest locus as the overall consensus of each cluster. The resulting loci with expanded boundaries and selected consensus were supplied to our pipeline for standard motif re-decomposition and efficient motif selection. Finally, we expanded the original vamos genomic catalog by adding loci that were completely uncovered in the catalog. Expansion for the CHM13 reference was done similarly by lifting over the TRExplorer v1.0 GRCh38 catalog onto CHM13 before loci merging and consensus refinement.

#### 2. Tandem repeat analysis using vamos output (tryvamos)

The variation annotated by vamos differs from conventional sequence variants because they are highly multiallelic and differ in both variant length and sequence composition. We developed a toolset tryvamos that has functions to analyze vamos TR annotations. This toolset is designed to accept vamos VCF files as input and provides functions for panel VCF combination (--combineVCF), feature matrix generation for high-dimensional analysis (--quickFeature), panel allele visualization (--waterfallPlot) (Supplementary Figure S3-5), allelic comparison of paired samples (--pairwiseCompare), as well as genome wide tests and visualization of divergent loci between two panels (--testTwoPanels). As the scope of LRS studies increases, the tryvamos toolset will undergo iterative enhancements to include new capabilities.

### Supplementary Figures

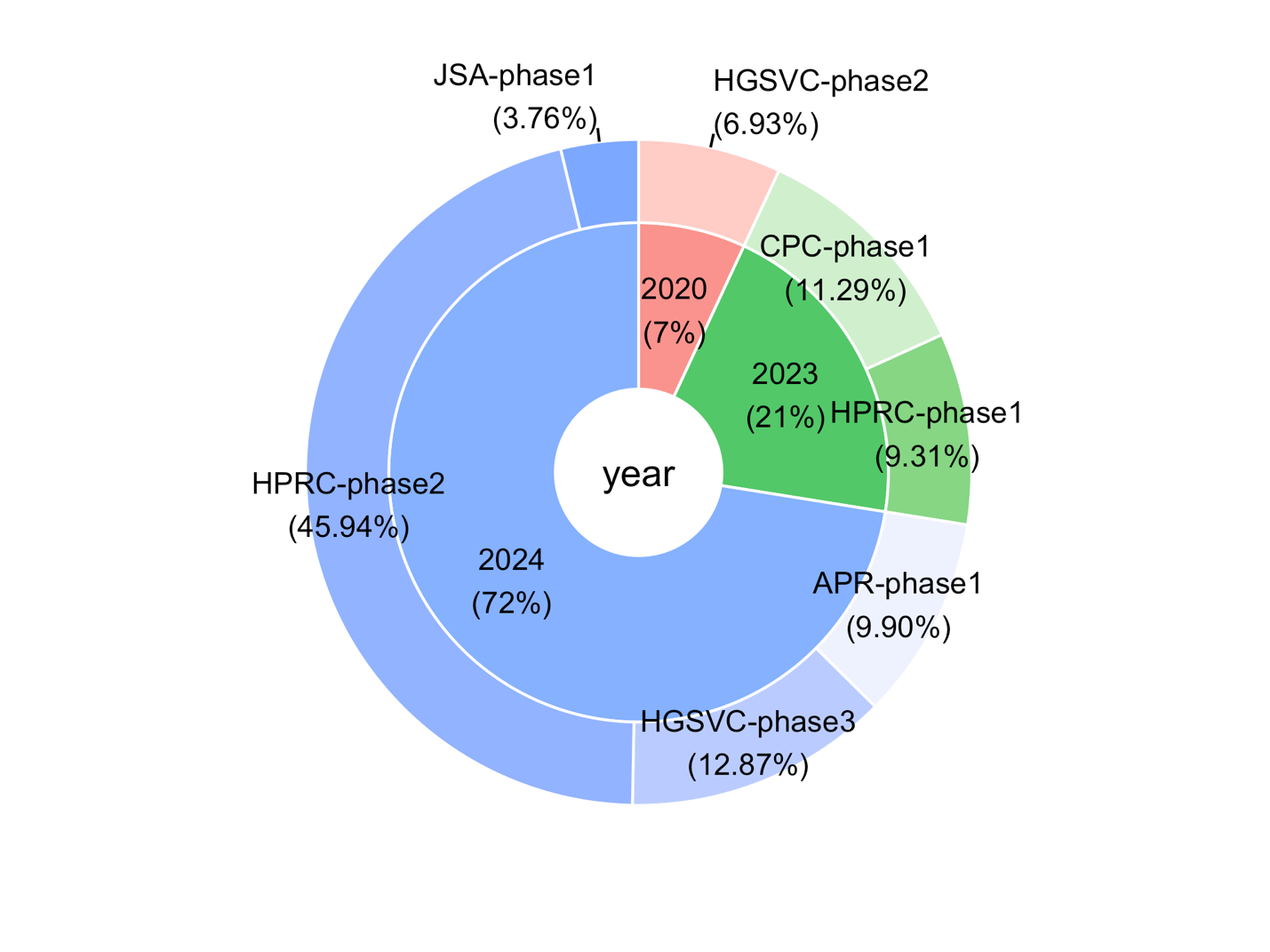

**Supplementary Figure S1**. Distribution of 505 diploid HiFi assemblies from seven sequencing consortia.

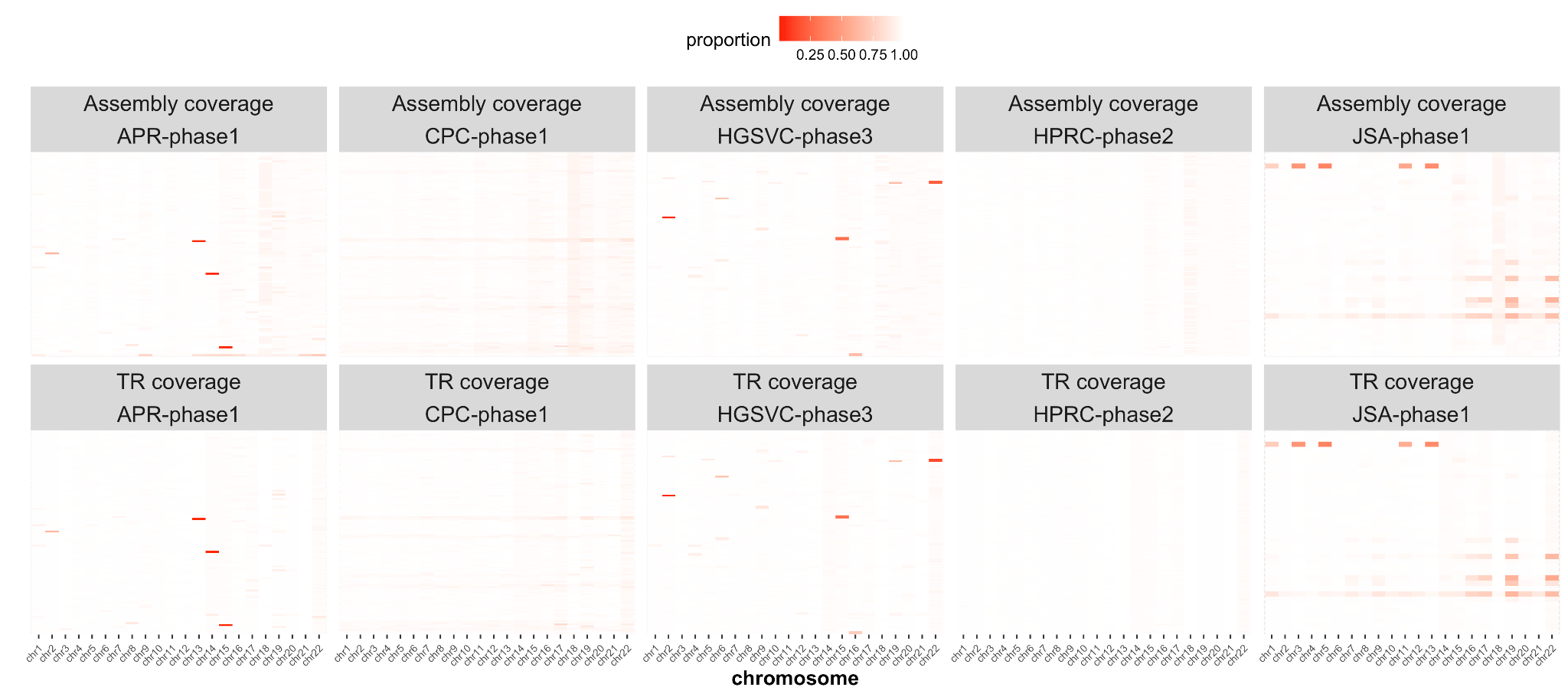

**Supplementary Figure S2**. Genomic coverage (upper panel) and TR coverage by vamos (lower panel) of genomes in 5 major HiFi sequencing consortia by genome and chromosome. Darker color indicates poorer coverage.

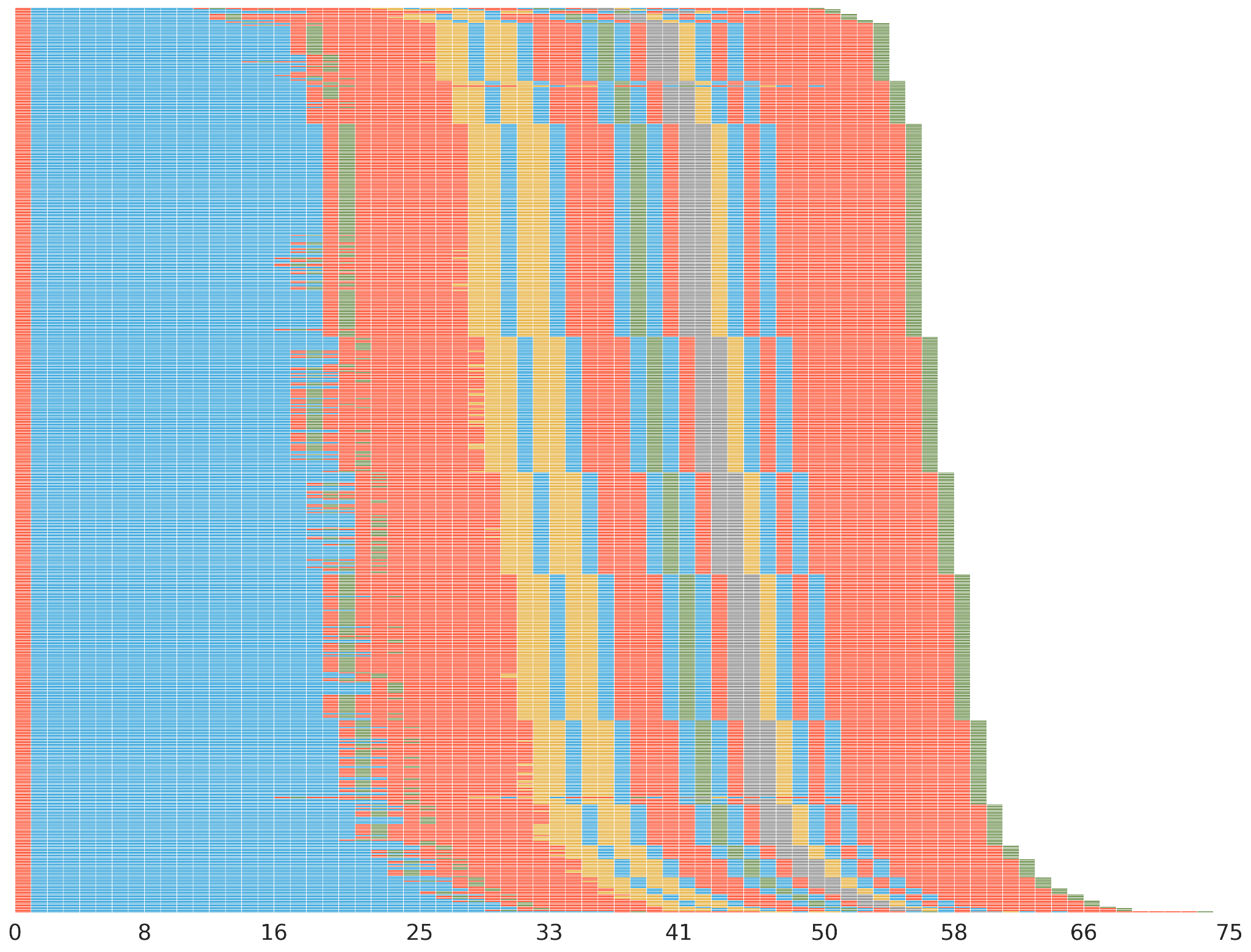

**Supplementary Figure S3**. Waterfall plot of the *HTT* coding STR before manual curation (chr4:3074876-3075052) on 594 HGSVC Phase 3 and HPRC Phase 2 genomes. The locus boundary includes the extra CCG repeats following the biologically recognized CAG repeats.

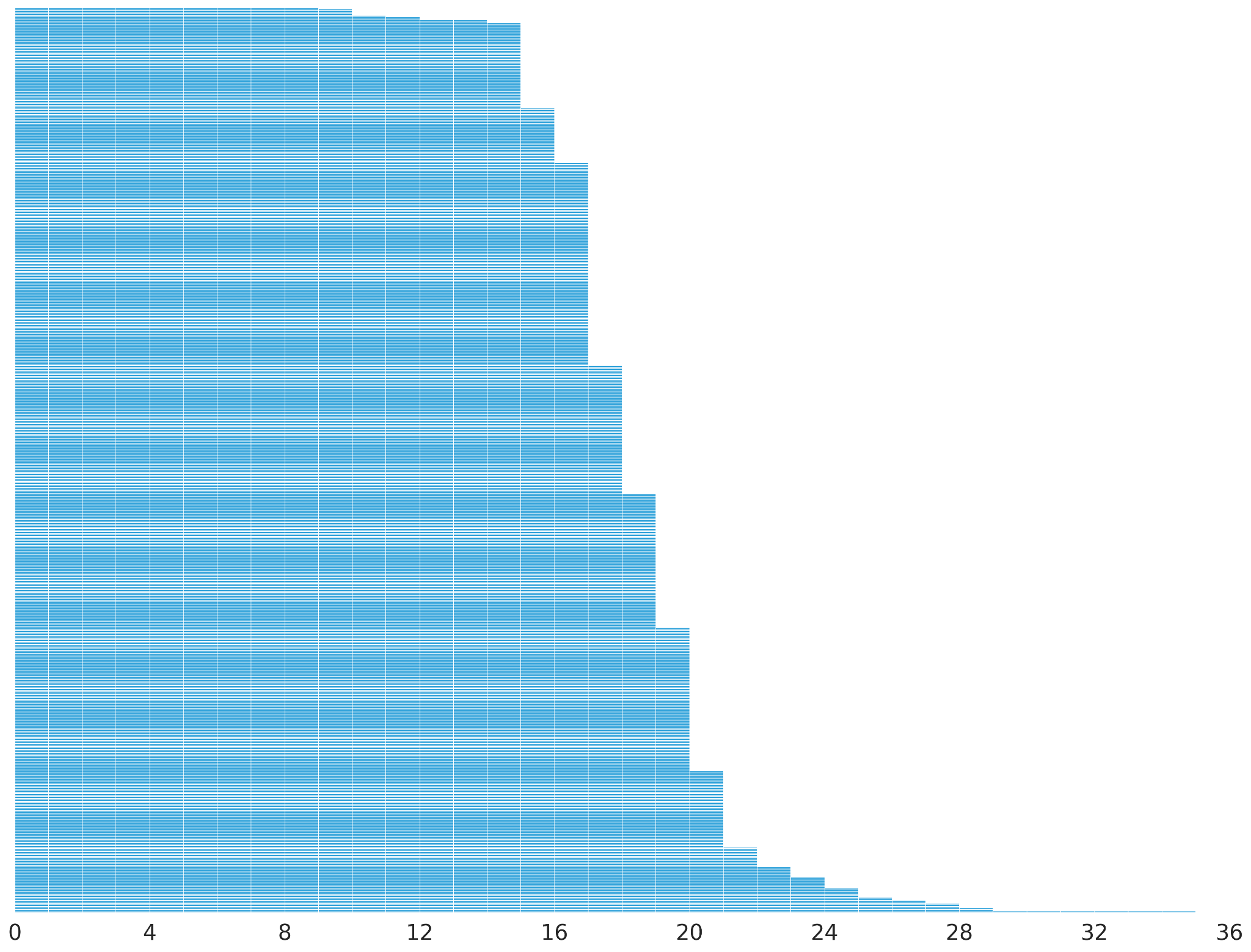

**Supplementary Figure S4**. Waterfall plot of the manually curated *HTT* coding STR (chr4:3074876-3074933) on 594 HGSVC Phase 3 and HPRC Phase 2 genomes. The locus boundary is curated to exclude the extra CCG repeats, leaving only the CAG repeats to reflect the biologically recognized allele length.

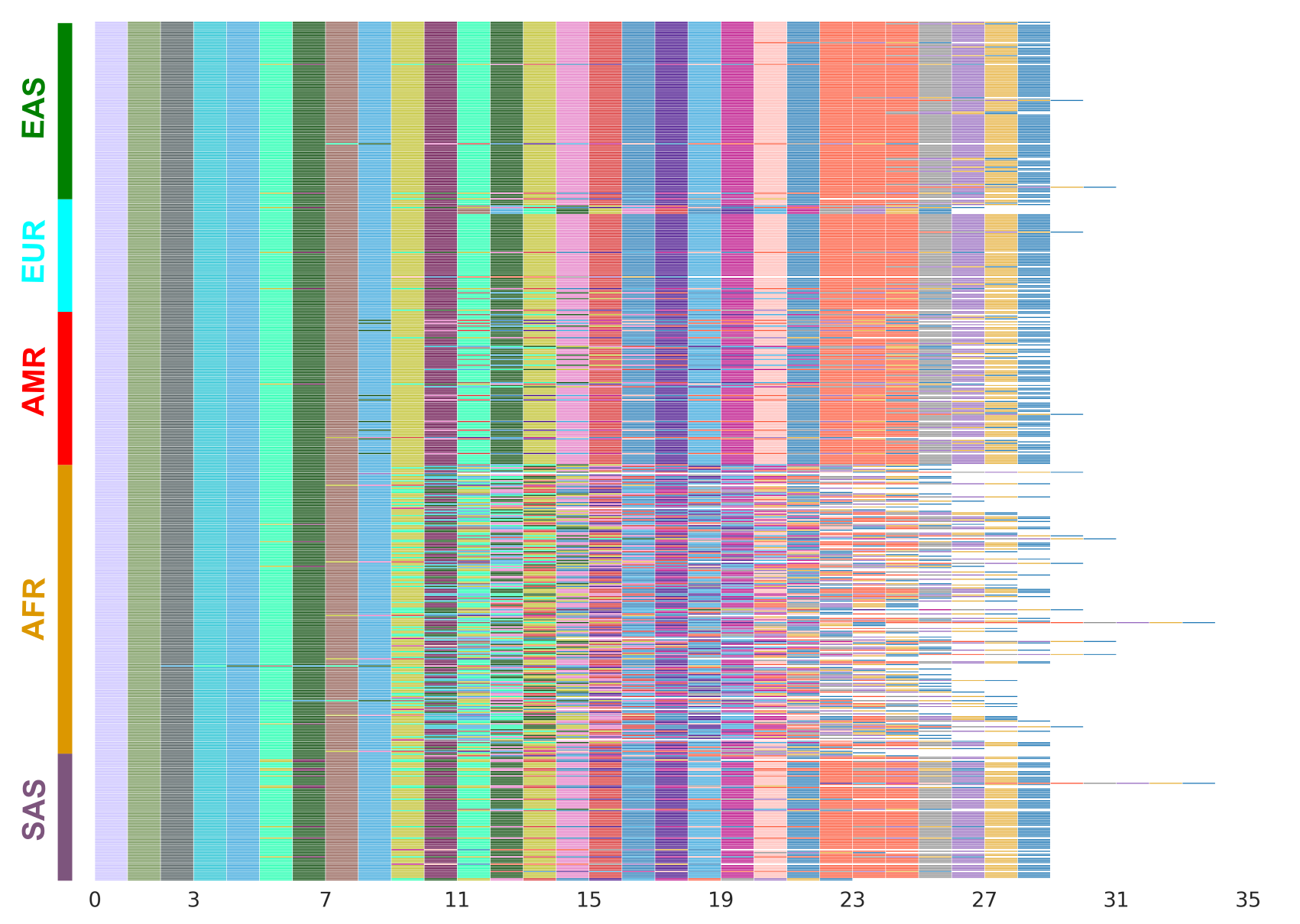

**Supplementary Figure S5**. Waterfall plot of the *PLIN4* coding VNTR (chr19:4510838-4513560) on 594 HGSVC Phase 3 and HPRC Phase 2 genomes. The African population exhibits the highest allele length and compositional diversity while the East Asian population exhibits the lowest allele length and compositional diversity.

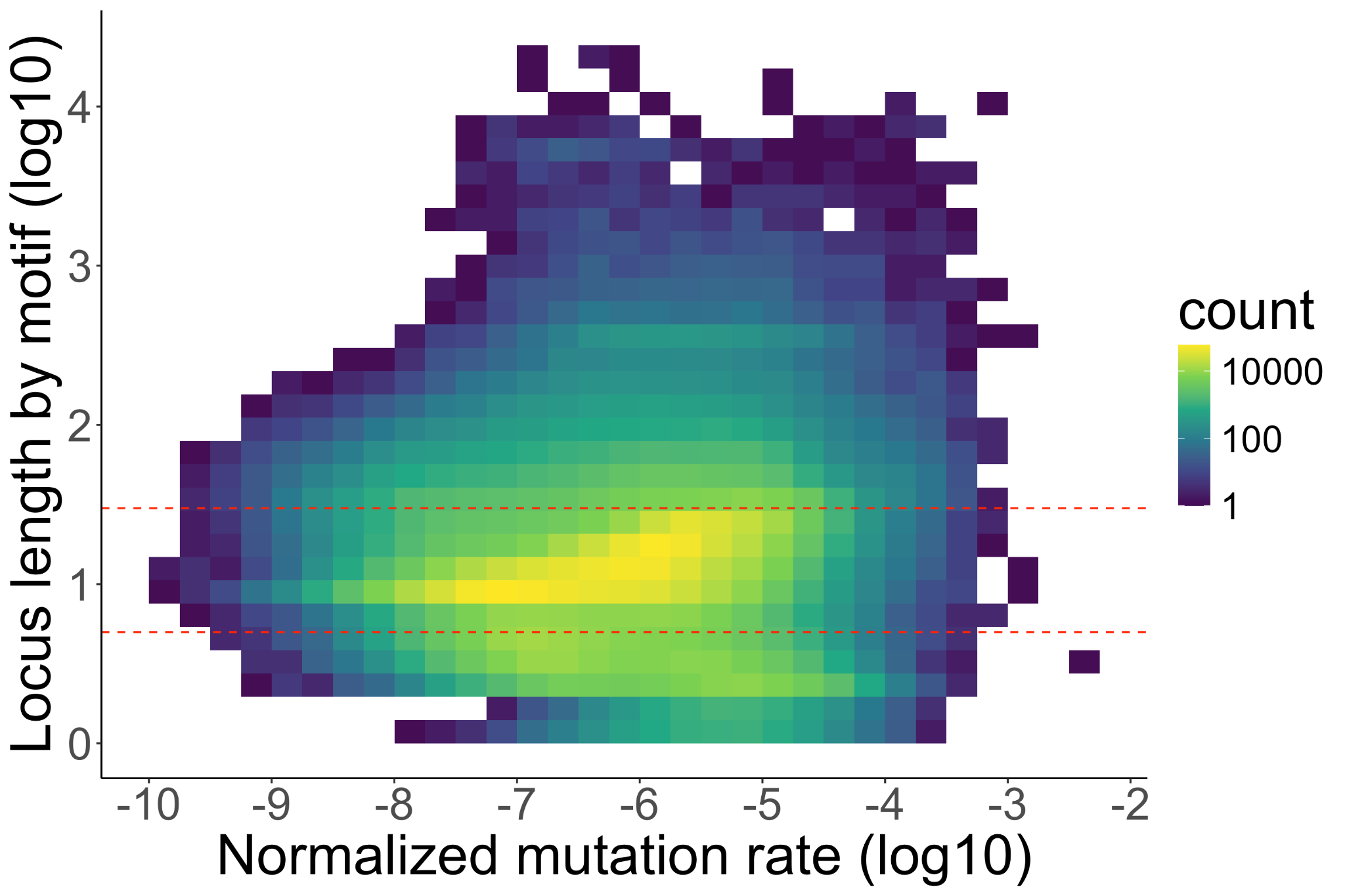

**Supplementary Figure S6**. Distribution of TR mutation rate by locus length. Both axes are on log10 scale. A linear trend is shown on loci whose average motif copy numbers are between 5 and 30 (marked by the red dashed lines). This implies a power relationship between motif copy number and mutation rate on these loci.

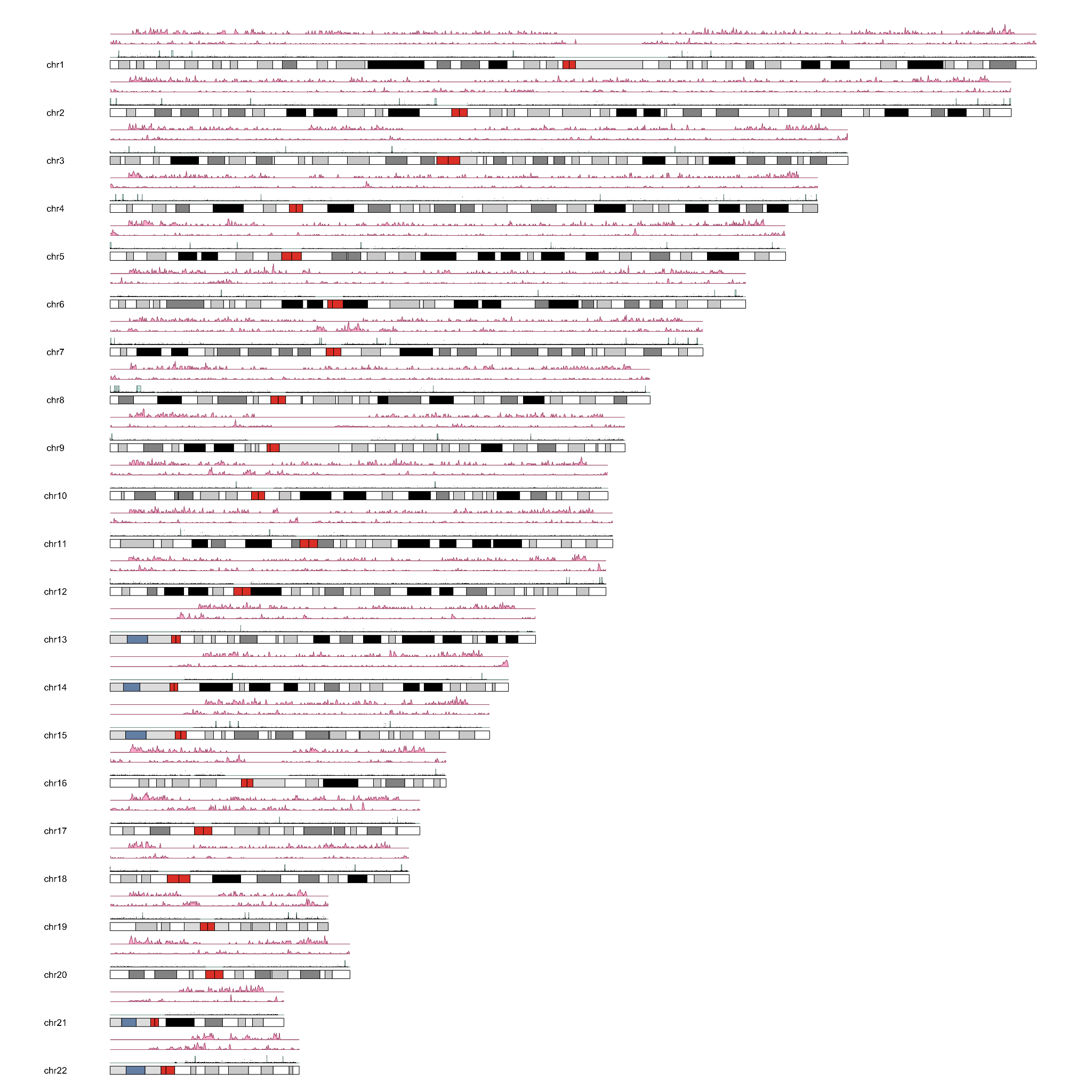

**Supplementary Figure S7**. TR mutational hotspots on GRCh38. Bottom track, mutation rate (black) and segmented hotspots (green). Middle track, distribution of segmental duplications. Upper track, distribution of recombination hotspots.

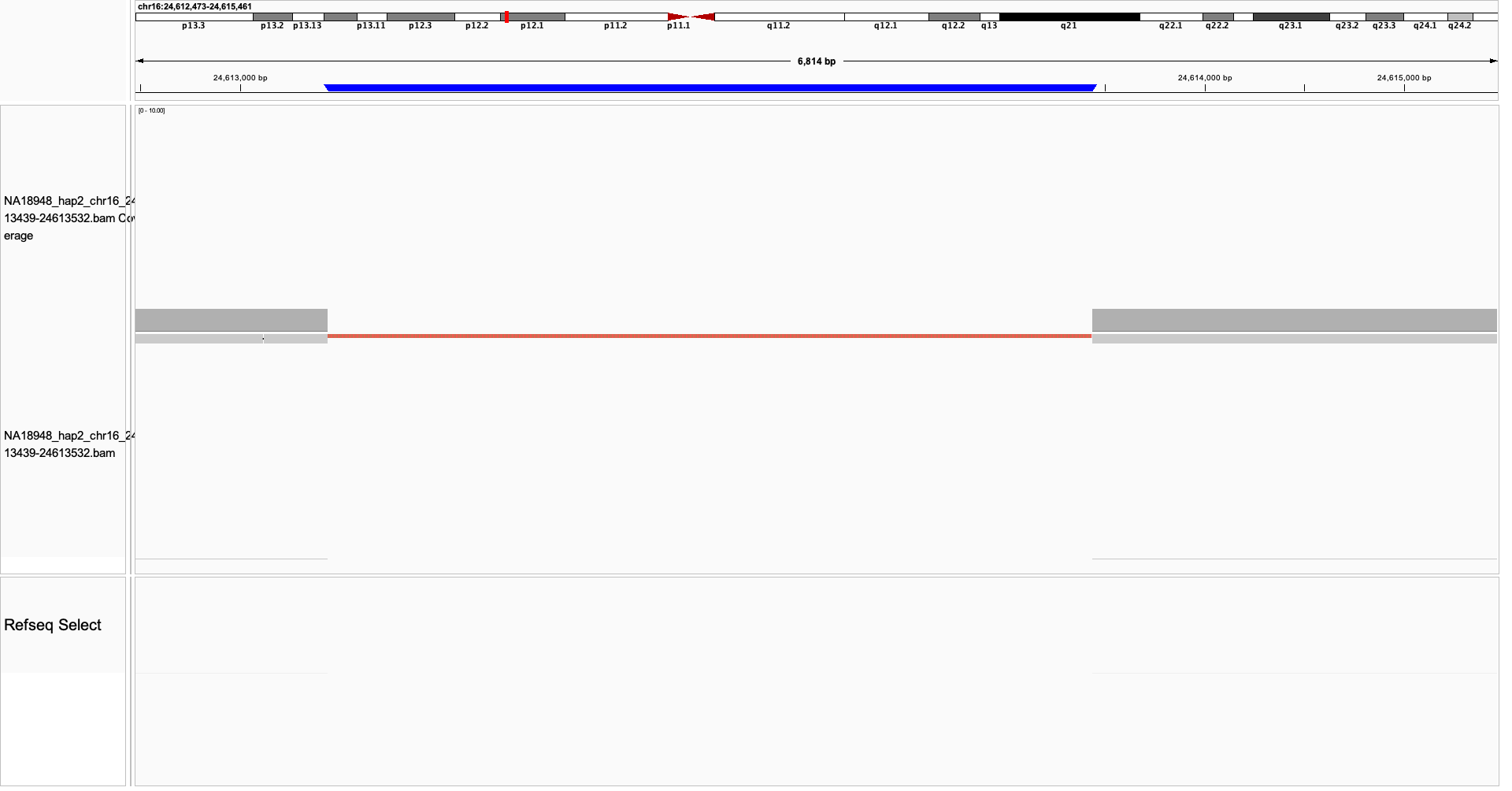

**Supplementary Figure S8**. IGV plot of an outlier allele of the ATTTT STR at chr16:24613439-24613532. The STR was reported to be associated with FAME6. The plotted genome (HPRC Phase 2 NA18948_hap2) has an insertion of 3,821bp of the ATTTT motif.

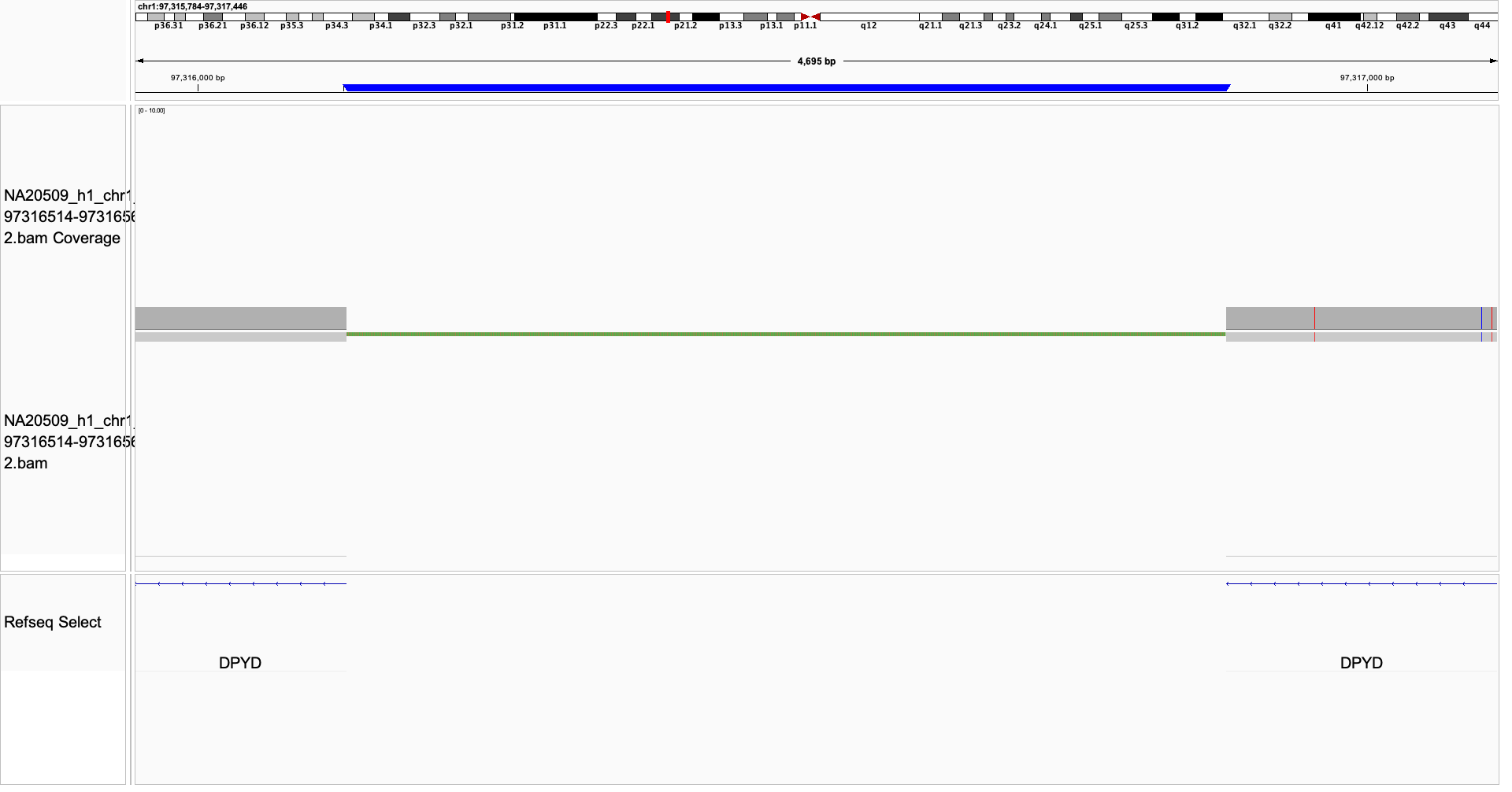

**Supplementary Figure S9**. IGV plot of an outlier allele of the AAATA STR at chr1:97316514-97316562. The plotted genome (HGSVC Phase 3 NA20509_h1) has an insertion of 3,029bp of the AAATA motif.

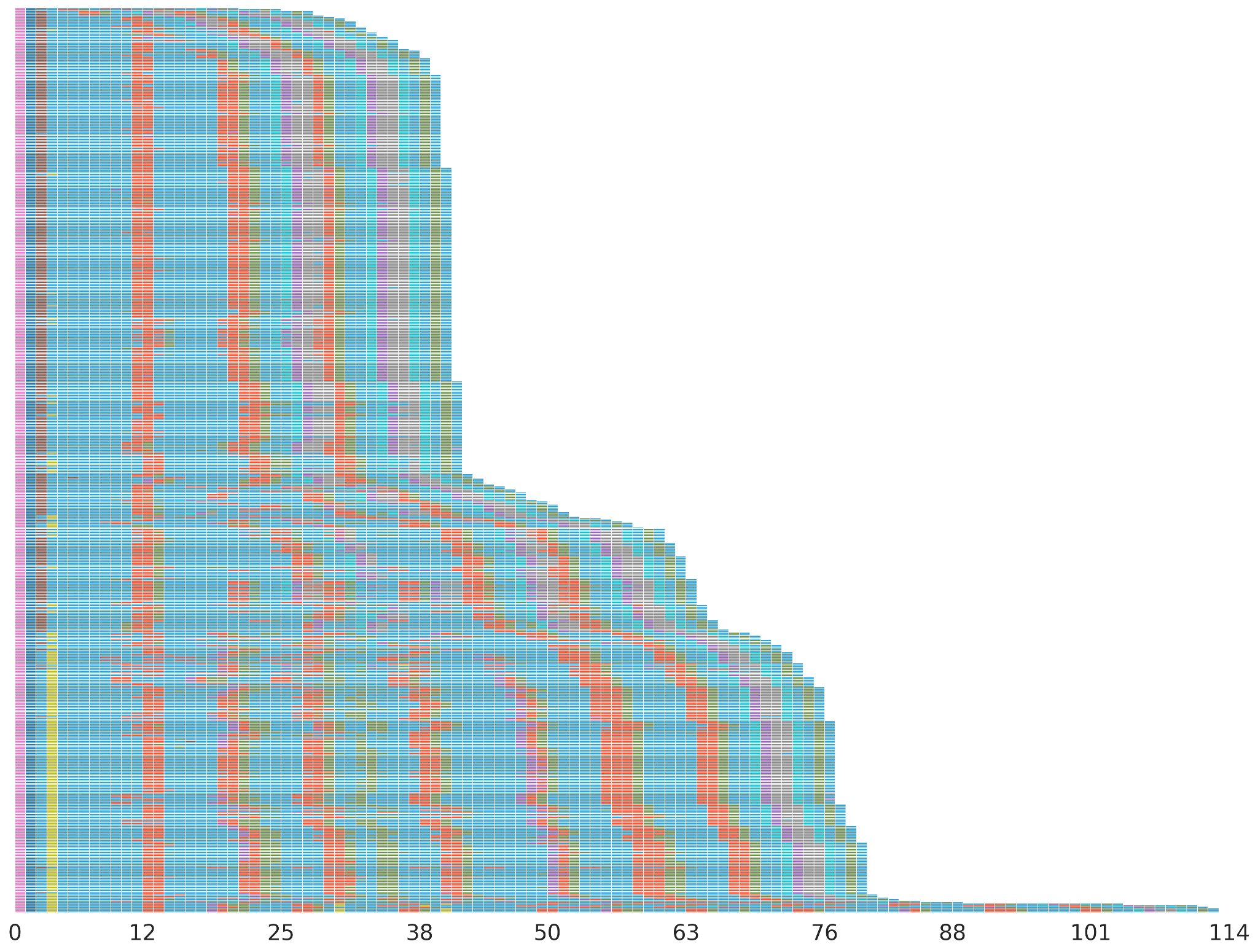

**Supplementary Figure S10**. Waterfall plot of the *MUC1* coding VNTR (chr1:155188482-155192051) on 594 HGSVC Phase 3 and HPRC Phase 2 genomes. Block mutations of multiple motifs form a higher order of repetitive patterns.

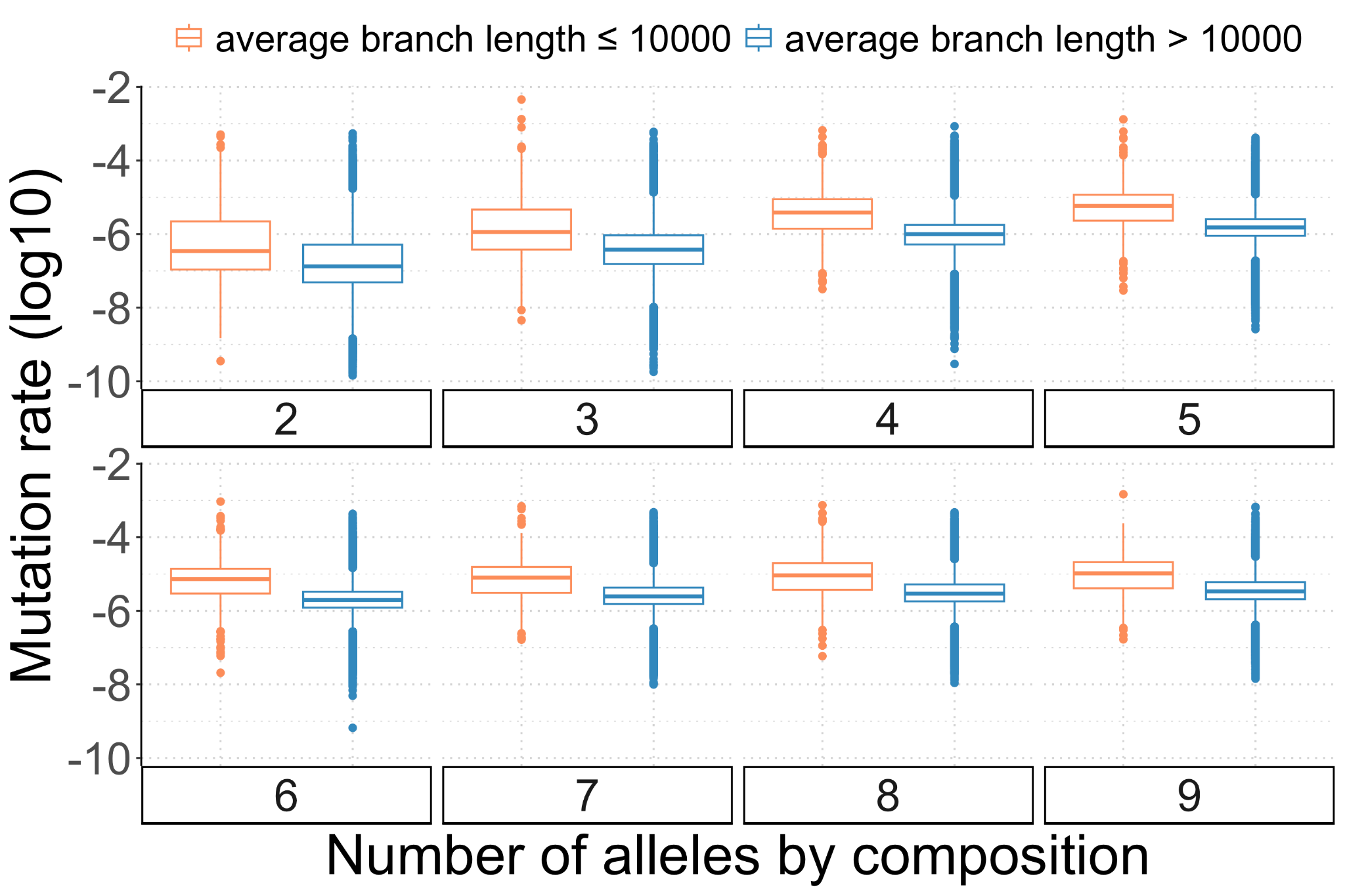

**Supplementary Figure S11**. Effect of tree branch length on estimated TR mutation rate by the number of TR alleles.

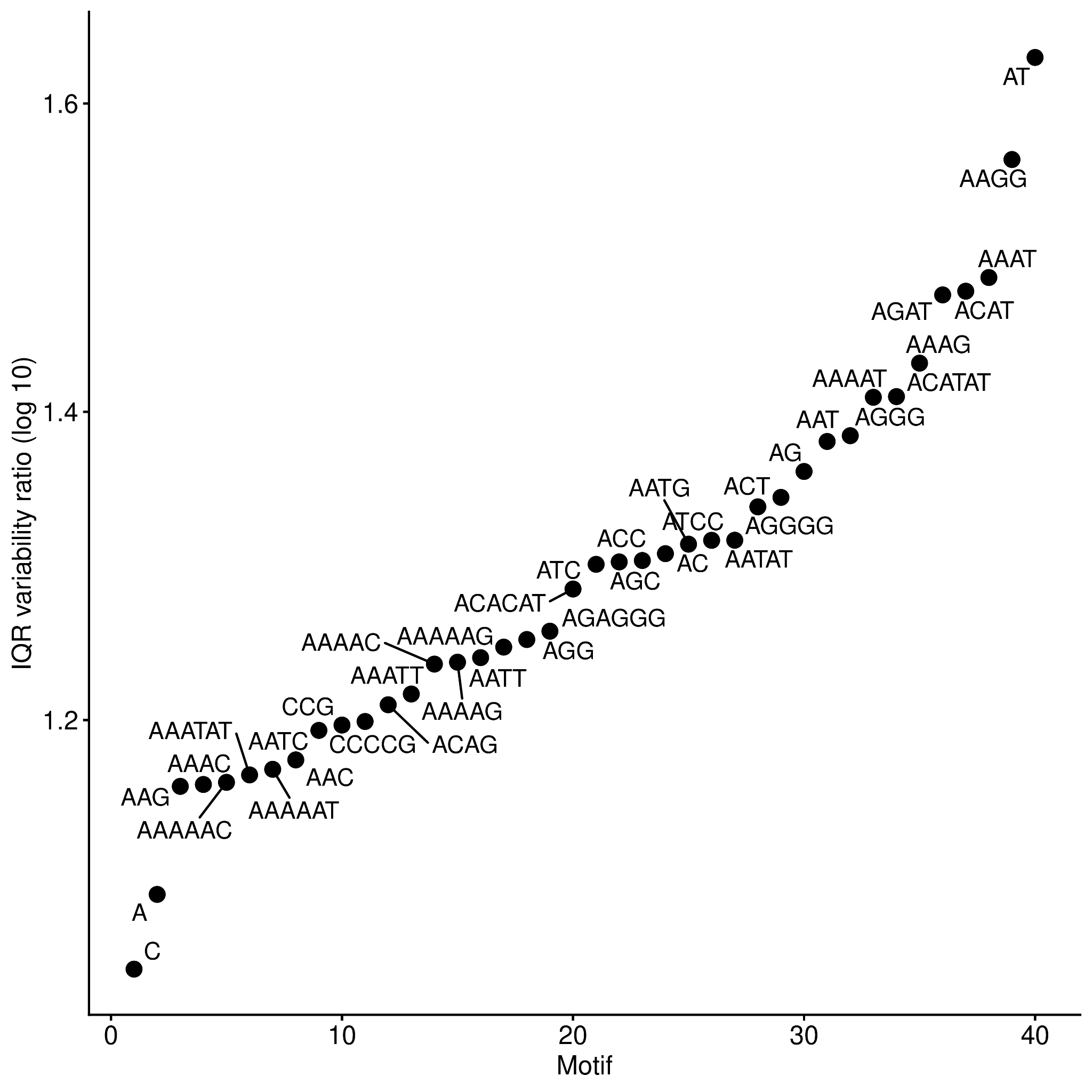

**Supplementary Figure S12**. Range of mutation rate by STR motif.

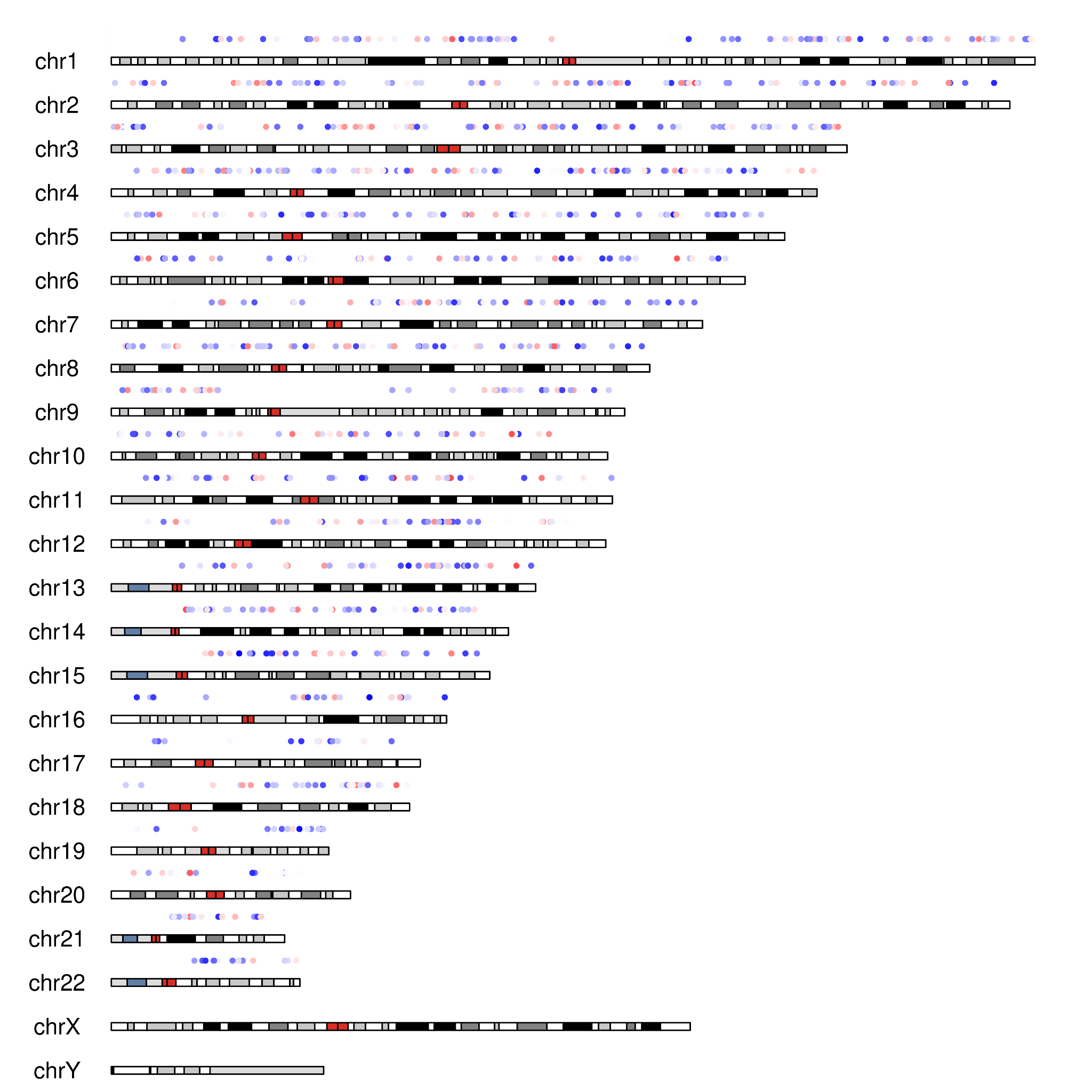

**Supplementary Figure S13**. Genome wide distribution of AT STR sequences colored by mutation rate for highest (red) to lowest (blue) for 1000 random sampled loci.

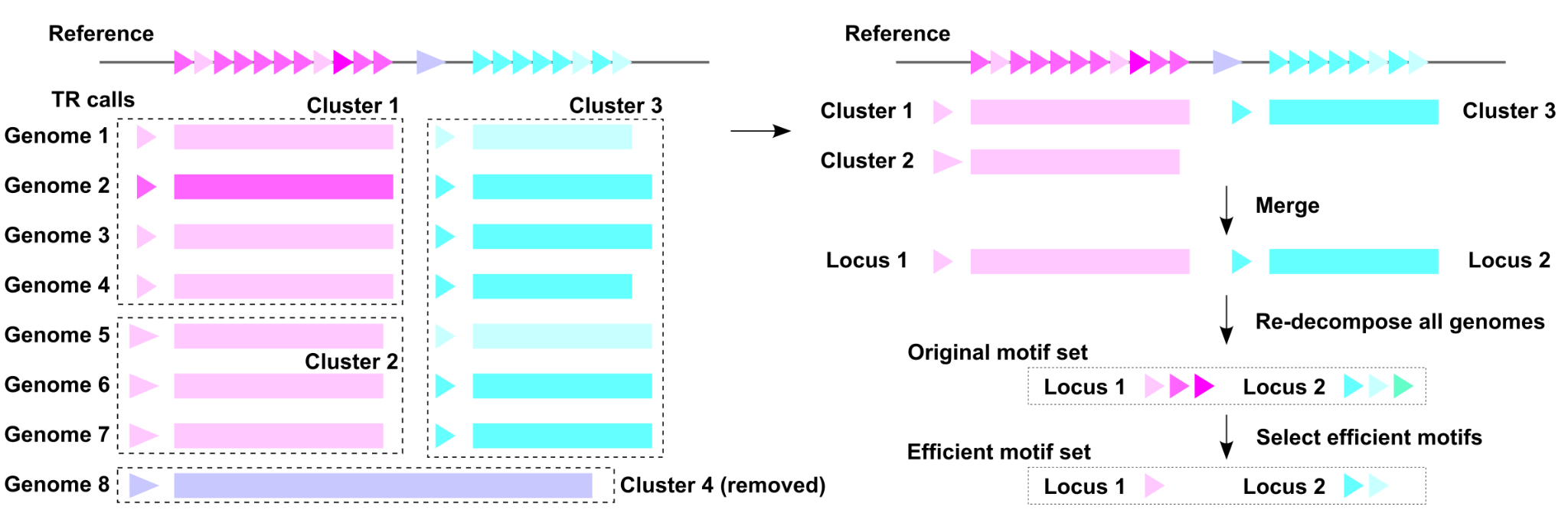

**Supplementary Figure S14**. Overview of the clustering algorithm for the construction of TR boundaries and motifs. Preprocessed TR calls (TRF and RepeatMasker) are first clustered by boundary coordinates and length of consensus motifs before merging across clusters. Poorly supported clusters are removed, and the consensus motif of the best supported cluster is selected for each locus after merging. Next, the motif set used for efficient motif selection is obtained by re-decomposing all input genomic TR sequences using the consensus motif, followed by selection of efficient motifs.
